# Chromatin-focused genetic and chemical screens identify BRPF1 as a targetable vulnerability in Taxol-resistant triple-negative breast cancer

**DOI:** 10.1101/2024.04.16.587277

**Authors:** Ozlem Yedier-Bayram, Ahmet Cingöz, Ebru Yilmaz, Ali Cenk Aksu, Beril Esin, Nareg Pınarbaşı-Değirmenci, Ayse Derya Cavga, Beyza Dedeoğlu, Buse Cevatemre, Hamzah Syed, Martin Philpott, Adam P. Cribbs, Udo Oppermann, Nathan A. Lack, Ceyda Acilan, Tamer T. Onder, Tugba Bagci-Onder

## Abstract

Triple-negative breast cancer (TNBC) stands out as a particularly aggressive and frequently recurring form of breast cancer. Due to the absence of hormone receptors, the available treatment avenues are constrained, making chemotherapy the primary approach. Unfortunately, the development of resistance to chemotherapy poses a significant challenge, further restricting the already limited therapeutic alternatives for recurrent cases. Understanding the molecular basis of chemotherapy resistance in TNBC is pivotal for improving treatment outcomes. Here, we generated two different Taxol-resistant TNBC cell lines with a dose-escalation method to mimic chemotherapy resistance *in vitro*. These cells exhibited hallmark features of resistance, including reduced cell growth, altered morphology, and evasion of apoptosis. Transcriptome analysis uncovered elevated *ABCB1* expression and multidrug-resistant phenotype in the resistant cells. To comprehensively investigate the key epigenetic regulators of Taxol resistance, we conducted chromatin-focused genetic and chemical screens and pinpointed Bromodomain and PHD Finger Containing 1 (BRPF1) as a novel regulator of Taxol resistance in TNBC cells. Knockout of BRPF1, the reader protein in the MOZ/MORF histone acetyl-transferase complex, but not the other complex members, sensitized resistant cells to Taxol. Additionally, BRPF1 inhibitors, PFI-4 and OF-1, in combination with Taxol significantly reduced cell viability. Transcriptome analysis upon BRPF1 loss or inhibition revealed a negative impact on ribosome biogenesis-related gene sets, resulting in a global decrease in protein translation in Taxol-resistant cells. Our ChIP-qPCR analysis demonstrated that active BRPF1 directly interacts with the *ABCB1* promoter, enhancing its expression towards inducing a multidrug-resistant phenotype. Conversely, knockout or inhibition of BRPF1 leads to decreased ABCB1 expression. This dual mechanism critically sensitizes Taxol-resistant TNBC cells to chemotherapy. Our findings uncover a comprehensive molecular framework, highlighting the pivotal role of epigenetic reader protein BRPF1 in Taxol resistance and providing potential avenues for therapeutic intervention in TNBC.

## INTRODUCTION

Triple-negative breast cancer (TNBC) is a very aggressive and recurrent type of breast cancer that predominantly affects younger women [1]. TNBC tumors are characterized by the absence of estrogen receptor (ER), progesterone receptor (PR), and human epidermal growth factor receptor-2 (HER2) overexpression. The lack of hormone receptor expressions limits targeted therapy options, leaving taxane and anthracycline-based chemotherapy as the mainstay treatments [2].

Taxol (Paclitaxel), a member of the taxane class, exerts its therapeutic effect by inducing mitotic catastrophe. It binds to and stabilizes microtubules, preventing their disassembly during metaphase, resulting in mitotic arrest at the G_2_/M checkpoint and eventual cell death [3, 4]. Although TNBC patients initially respond to taxane-based chemotherapy, they are more prone to developing resistance and recurrence than hormone receptor-positive counterparts [2, 5]. Taxol resistance is often linked to increased expression of ABC transporters, particularly ABCB1 (P-glycoprotein, MDR1), which actively pumps the drugs out of the cell. The broad substrate specificity of the ABC transporters limits the use of alternative chemotherapeutics once they are overexpressed, leading to a multidrug resistance phenotype (MDR) [6]. Efforts to overcome MDR through the combined use of ABC transporter inhibitors and chemotherapeutics have faced challenges [7]. First-generation ABCB1 inhibitors, such as verapamil, resulted in toxic side effects without additional benefits [8]. Although second-generation inhibitors aimed to enhance specificity and reduce side effects, they inadvertently increased systemic exposure, exacerbating toxicity. Third-generation ABCB1 inhibitors, despite effectively reversing ABCB1-mediated MDR, did not improve overall survival rates across various cancer types [9–11]. Moreover, the possibility of compensation through the co-expression of different ABC transporter family members complicates targeted inhibition strategies. Understanding the upstream regulators specific to cancer for ABC transporters becomes essential due to the toxicity linked with direct inhibition of ABCB1 and the potential for compensatory actions by other transporters.

Advances in single-cell sequencing technologies have revealed non-genetic mechanisms and pre-existing epigenetically poised cells that contribute to intra-tumor heterogeneity and clonal selection during treatment [12–16]. For instance, heterogeneity in histone modifications, such as H3K27me3, has been linked to the emergence of Taxol-resistant subpopulations [17]. EZH2 inhibitors that decrease histone methylation sensitized resistant cells to paclitaxel and reduced metastasis [17–19]. Conversely, it has been shown that H3K27me3 prevents cells from escaping chemotherapy, acting as a lock in drug-tolerant persister cell-specific genes in TNBC cells [20]. Inhibitors of H3K27me3 demethylases, combined with 5-FU, reduced the number of drug-tolerant cells [20]. Pre-existence of drug-tolerant tumor cells in populations with increased levels of KDM5A and hypersensitivity to HDAC inhibitors has also been shown [21]. Chromatin remodeling factors regulating the accessibility of stemness genes in TNBC contribute to invasive tumors [22]. Alternatively, chromatin regulators such as BAF and COMPASS complexes and KDM4B have been found to increase anthracycline sensitivity by enhancing chromatin accessibility [23]. Collectively, these findings highlight the significant impact of epigenetic regulators on the development of chemoresistance, while their specific roles might be tissue-, cancer-, and time-dependent.

The link between epigenetic regulation of Taxol resistance and the MDR phenotype remains poorly defined. Earlier studies showed that loss of repressive marks on the *ABCB1* promoter, such as DNA methylation, leads to increased expression of ABCB1 in different cancer types [24–27]. A correlation between increased KDM5A levels and ABCB1 expression was demonstrated in lung adenocarcinoma [28]. On the other hand, a recent study proposed that higher-order 3D genome topology is crucial for *ABCB1* activation [29]. While modulation of DNA methylation or histone acetylation had no clear impact on ABCB1 expression, dissociation of the ABCB1 locus from the nuclear lamina led to its activation in Taxol-resistant cells. These data underscore the absence of a clear consensus on how ABCB1 is epigenetically regulated, emphasizing the need for further investigation.

In this study, we examined the epigenetic modifiers regulating Taxol resistance in TNBC cells by first generating Taxol-resistant cells *in vitro*. Characterization of resistant cells revealed that *ABCB1* was highly upregulated. Using our chromatin-focused CRISPR/Cas9 library, EPIKOL [30], and epigenetic probe-library concurrently, we identified epigenetic vulnerabilities in Taxol-resistant cells. Bromodomain and PHD Finger Containing 1 (BRPF1) protein, a member of a histone acetyltransferase complex, emerged as a regulator of Taxol-resistance and ABCB1 expression. Transcriptome analysis also demonstrated that BRPF1 contributes to elevated ABCB1 protein levels and regulates ribosome biogenesis, impacting protein translation. The combination of BRPF1 inhibitors with chemotherapy for ABCB1-high tumors presents a promising therapeutic opportunity.

## METHODS

### Cell culture

SUM159PT TNBC cell line and HEK293T cells were kind gifts from Robert Weinberg (MIT, Boston, USA). HEK293T cells were cultured in DMEM (Gibco, USA) supplemented with 10% fetal bovine serum (Gibco, USA) and 1% penicillin/streptomycin (Gibco, USA). SUM159PT cells were cultured in Ham’s F12 nutrient mix (Gibco, USA) supplemented with 5% FBS, 5 µg/ml insulin (Sigma-Aldrich, USA), 1 µg/ml hydrocortisone (Sigma-Aldrich, USA) and 10 mM HEPES (Thermo Fisher, USA). Cells were maintained in a humidified incubator at 37°C with a 5% CO_2_ level. All cell lines were tested regularly for mycoplasma infection.

### Cell viability assay and determination of IC_50_ values

Cells were seeded into 96-well black plates as 2000 cells/well for SUM159PT cells and its derivatives. Next day, cells were treated with corresponding chemicals. For IC_50_ determination, Taxol (Paclitaxel, Sigma) was serially diluted 3.16-fold (starting from 10 µM to 0.1 nM). After 72 hours of treatment with the indicated chemical, media were discarded, and a luminescence-based cell viability assay (CellTiter-Glo®, Promega, USA) was performed according to the manufacturer’s recommendations. Results were analyzed using GraphPad Prism 8.

### Generation of drug-resistant TNBC cell lines

Cells were seeded as 2x10^5^ cells/well in 6-well plates. Next day, they were treated with either IC_10_ (0.4 nM) or IC_50_ (1.5 nM) values of Taxol for 72 hours as starting concentrations. When cells became confluent under drug treatment, concentrations were doubled. If cells did not look healthy or were too sparse, they were taken into drug-free medium until they form colonies. Upon reaching confluence, the last concentration of drug was applied again, and the procedure was repeated. Alongside, DMSO-treated parental cell lines were aged as controls. Drug treatment was continued until a significant difference between IC_50_ values of parental and resistant cell lines was obtained. Resistant cells derived from SUM159PT cells were named according to the initial and final doses of Taxol applied. Cells were maintained at the final dose of Taxol (160 nM for T1-160 and 450 nM for T2-450) while culturing.

### Colony formation assay

Cells were seeded as 1000 cells/well for parental and 1500 cells/well for resistant cells in triplicate in 6-well plates. For sgRNA-containing experiments, cells were seeded after puromycin selection. Next day, drug treatments were performed as indicated in each experiment. 72 hours later, medium was refreshed, and cells were allowed to grow for 10-14 days. At the end of the incubation period, media were discarded, cells were washed with PBS and fixed with ice-cold 100% methanol for 5 minutes. Methanol was discarded and cells were stained with crystal violet for 15 minutes. Quantification of the area occupied by colonies was performed by using ImageJ [31].

### Annexin V Staining

Annexin V staining was performed with the Muse® Annexin V & Dead Cell Kit (Luminex, MCH100105) according to the manufacturer’s instructions. Briefly, 75.000 cells were collected with their media and centrifuged at 1200 rpm for 5 minutes. After washing with 500 µl of cold PBS with 1% FBS, pellet was resuspended in 75 µl of cold PBS with 1% FBS and mixed with 75 µl of Annexin V & Dead Cell Reagent. Samples were incubated at room temperature for 20 minutes and analyzed with Muse Cell Analyzer (Merck, Darmstadt, Germany) with 5000 events per sample. Gates were determined according to parental cells.

### Western Blotting

Cells and the media were harvested and centrifuged at 10.000 rpm at 4°C for 5 min. Pellet was dissolved in NP-40 lysis buffer in an appropriate volume (50mM Tris Buffer pH 7.4, 250mM NaCl, 5mM EDTA, 50mM NaF, 1%NP40, 0.02% NaN3, 1mM PMSF).

Protein concentration was determined by Pierce’s BCA Protein Assay Kit (Thermo Scientific, 23225, USA). 50µg protein for each sample was mixed with 4X Laemni sample buffer (Biorad, 1610747, USA) containing 10% beta-mercaptoethanol and incubated at 95°C for 10 minutes. Samples were loaded and run on a 4-12% Mini Protean TGX Precast Gel (Biorad, 456-1044, USA). Proteins were transferred onto PVDF membranes via Bio-RadTrans-Blot® Turbo™ Transfer System (Bio-Rad, USA). The membrane was blocked with 5% non-fat dry milk in TBS-T for one hour at room temperature. Then, the membrane was incubated with primary antibody overnight on a shaker at 4⁰C. Primary antibodies used in this study are listed in **Supplementary Table 1**. After washing with TBS-T, membranes were incubated with secondary antibodies conjugated to HRP diluted 1:10000 in TBS with 5% non-fat dry milk for 1 hour at room temperature. Signals were detected with either with SuperSignal™ West Femto Maximum Sensitivity Substrate (Thermo Scientific, 34095, USA) or ECL in ChemiDoc XRS+ System (Biorad, USA).

### Cell Cycle Analysis

1x10^6^ cells were collected and washed with PBS. Pellet was resuspended in 100 µl PBS. Cell suspension was added drop by drop into the freshly prepared, cold 1 ml of 70% Ethanol while vortexing for fixation. Samples were kept at -20°C for 24 hours. 200 µl of fixed cells were washed with PBS twice and resuspended in 150 µl of Muse Cell Cycle Reagent (The Muse® Cell Cycle Kit (Luminex, MCH100106)). Samples were incubated with the reagent at room temperature for 30 minutes in the dark and run through the Muse Cell Analyzer (Merck, Darmstadt, Germany) with 10.000 events per sample and analyzed with the Muse Cell Analyzer software. Gates were determined according to parental cells.

### Immunofluorescence

Cells were fixed with ice-cold 100% methanol for 10 minutes at -20°C. After washing with PBS twice, cells were permeabilized with 0.01% Triton X-100 at room temperature for 5 minutes. Cells were blocked with SuperBlock (ScyTek laboratories, Logan, UT, USA) for 15 minutes at room temperature and incubated with the primary antibody anti-α-tubulin (DM1A) (Sigma T9026) 1:10000 diluted in SuperBlock for 1 hour at room temperature. After washing, cells were incubated with Alexa Fluor 488 Anti-mouse secondary antibody (Invitrogen) for 1 hour at room temperature in the dark, and coverslips were mounted with DAPI (Mounting medium with DAPI, Abcam, #ab104139). Images were taken using Zeiss Axio Imager M1 (Germany) at 40x magnification.

### RNA sequencing and transcriptome analysis

Total RNAs were isolated by using MN Nucleospin RNA isolation kit according to manufacturer’s instructions. Library preparation and sequencing were performed at University of Oxford (Oxford, UK). Briefly, RNA was DNase I-treated, cleaned, and concentrated (Zymo RNA Clean and Concentrator, Zymo Research), then enriched for poly(A) mRNA (NEBNext poly(A) mRNA Magnetic Isolation Module, NEB Biosystems, Ipswich, UK). Sequencing libraries were prepared using the NEBNext Ultra II RNA Library Prep kit (New England Biolabs). RNA quality was assessed using High Sensitivity RNA Screentape and an Agilent 4200 tapestation. Single-indexed and multiplexed samples were run on an Illumina Next Seq 500 sequencer using a NextSeq 500 v2 kit (FC-404–2005; Illumina, San Diego, CA) for paired-end sequencing. Bioinformatic analysis of the samples was performed to identify differentially expressed genes after pre-processing of the sequencing data. Reads were trimmed using trimmomatic [32], pseaudoaligned using kallisto [33] with an index built from the hg38 cDNA FASTA reference sequence, and then quality of the pseudoalignment was assessed using FastQC. Differential gene expression was performed using the DESEq2 package [34]. For parental-resistant comparisons, differentially expressed genes (DEGs) were defined with a threshold for Log_2_FoldChange>2 (up-regulated) or Log_2_FoldChange<-2 (down-regulated) and p<0.001. For drug or sgRNA-treated T1-160 cells, DEGs were determined as the genes that have padj<0.05. Gene set enrichment analysis was performed with Log_2_FoldChange rank-ordered gene lists by using GSEA software and all available gene sets from MsigDB at the date of January 24^th^, 2022 [35].

### Quantitative RT-PCR

RNA isolation and cDNA synthesis were performed as described [30]. List of qPCR primers can be found in **Supplementary Table 2**.

### Epigenetic Probe Library Screen

Chemical probe library targeting a wide range of epigenetic modifiers consists of 117 drugs (**Supplementary Table 3**). Library was a kind gift from Dr. Udo Oppermann, Oxford University, and was constructed as described [36]. Cells were seeded as 750 cells/well in 384-well plates. Next day, cells were treated with epigenetic probes alone or in combination with Taxol in a final concentration of 2.5 nM for Parental, 300 nM for T1-160 and 500 nM for T2-450. The Taxol doses were selected to correspond to IC_40_-IC_50_ in 384-well plate format. Epigenetic probes were added at 1:1000 dilution in triplicate as in the final concentrations indicated in **Supplementary Table 3.** 72 hours later, cell viability was measured by using CellTiter-Glo, and results were normalized to untreated controls. The mean cell viability and standard deviations (SD) of DMSO controls (in combination with Taxol) were calculated as 79.67% ± 0.7% for parental, 54.24% ± 1.43% for T1-160, and 62.20% ± 1.02% for T2-450 screens. An epigenetic probe was considered as a ‘hit’ if it decreased cell viability by 3 SDs of DMSO or lower in combination with Taxol (77.57% for parental, 49.95% for T1-160, and 59.15% for T2-450). We excluded the hits and refrained from labeling them on the plot if their negative control also affects cell viability.

### Chromatin-focused Knockout Screen

Epigenetic Knockout Library (EPIKOL) that targets 779 genes, including several controls, was used, and viruses were generated as described [30]. After Cas9-expressing T1-160 cells were generated, pooled lentiviral EPIKOL was transduced as 1000x representation with an MOI of 0.4. Transduced cells were selected with 5 µg/ml puromycin in the presence of 20 µM verapamil for 4 days, and 8 million cells were collected as the initial timepoint. Cells were divided into two groups and treated with either DMSO as a vehicle or 160 nM Taxol until they reached 16 population doublings. At the end of the culturing period, final cell pellets were collected as 8 million. Genomic DNA was isolated using Nucleospin Tissue kit (Macherey-Nagel, Germany) according to manufacturer’s instructions. Next generation sequencing (NGS) libraries were prepared as described [30]. Briefly, in the first PCR, 13.2 µg of gDNA was used as a template to achieve 250x coverage of the sgRNA library. In the second round, NGS (Illumina) adapters (stagger, index, and index read primer sequences) were added at the ends of PCR amplicons. Products were run on 2% agarose gel and purified using MN Nucleospin Gel and PCR Cleanup kit. NGS was performed at Genewiz (NJ, USA) by using Hiseq (Illumina) with at least 10 million reads/sample. Sequencing reads from R1 fastq files were aligned to the sgRNA library and counted using MAGeCK (v0.5.8). Biological replicates were presented as individual input files for counting, and results were normalized as Read Per Million (RPM) and converted to Log_2_ values. Counts of sgRNAs were then combined with median normalization to obtain gene-level log_2_ fold changes. p<0.05 cutoff was applied to gene-level analysis to identify significantly depleted genes.

### Dual-color competition assays

For validation of EPIKOL screen candidate hits, dual color competition assays were performed as described [30]. Briefly, T1-160-Cas9 cells expressing either PGK-H2BmCherry (Addgene #21217) or PGK-H2BeGFP (Addgene #21210) were seeded as 50.000 cells/well in 12-well plates. mCherry+ cells were transduced with LentiGuide-NT1 viruses, while eGFP+ cells were transduced with viruses carrying sgRNA of interest. 2 different sgRNAs were used per gene. List of sgRNAs can be found in **Supplementary Table 4**. Following puromycin selection, mCherry+ and eGFP+ cells were mixed in a 1:1 ratio and re-seeded into 24-well plates in two groups as triplicates. One day after seeding, Day0 measurements were taken by acquiring 3x3 images with a 4x objective in Cytation5 (BioTek, USA). Then, one group was treated with 160 nM Taxol, whereas the other was treated with DMSO as a vehicle. Cells were incubated for 24 days, with images taken on days 4,8,12,16, and 24. Number of mCherry+ and eGFP+ cells were counted from images using Gen5 software (BioTek, USA), and each measurement was normalized to Day0 to determine the percentage of GFP-positive cells.

### siRNA experiments

T1-160 cells were seeded as 300.000 cells/well in 6-well plates. Next day, Lipofectamine 3000 (Invitrogen, USA) transfection was performed according to manufacturer’s instructions. Briefly, one mixture containing 125 µl optimem with 7.5 µl Lipofectamine 3000 Reagent and another mixture containing 125 µl optimem with 100 pmol of either BRPF1 siRNA (Thermofischer Scientific, Cat no: 4392421, siRNA ID: s15422) or non-targeting siRNA (siNT) were prepared and vortexed well. siRNA mixture was added onto Lipofectamine 3000 mixture and incubated for 15 minutes. Final mixture was added to the cells dropwise, and 8 hours later, fresh media were given. Next day, cells were trypsinized and seeded for cell viability or colony formation assays.

### Functional Rescue Experiment

For overexpression of BRPF1 on cells carrying BRPF1 sgRNA, BRPF1 was cloned into a lentiviral vector, and the PAM sequence adjacent to the sgRNA binding site was mutated. For this, GFP-BRPF1 plasmid (Addgene #65382) was used as a template and amplified by using forward primers containing SalI and reverse primers containing XbaI cut sites. Resulting PCR product was cut and ligated to pENTR1A (Addgene #17398) entry vector. Then, LR reaction was performed to clone BRPF1 into pLEX_305-C-dTAG (Addgene #91798). PAM sequence adjacent to the BRPF1 sgRNA #1 was changed from NGG to NAG to ensure that the overexpression construct will not be knocked out by CRISPR/Cas9. Later, T1-160 cells were infected with BRPF1 sgRNA and the overexpression construct at the same time. On post-transduction day 5, cells were seeded for clonogenic assay.

### SUnSET Assay

To measure global translation rate, SUnSET assay, which non-radioactively measures puromycin labeled peptide amounts in a given period, was utilized. For this, T1-160 cells were infected with NT1 or BRPF1-targeting sgRNAs in the LentiGuide backbone with hygromycin selection. On day 10 after transduction, T1-160-NT1 cells were treated with Cycloheximide (5 µM) as a positive control for 48 hours. On day 12, T1-160 cells carrying NT1 or BRPF1-targeting sgRNAs were treated with puromycin (50 µg/ml) in the presence of verapamil (20 µM) for 30 minutes. Cells were scraped and stored at -80 °C.

### ChIP-qPCR

For Chromatin immunoprecipitation (ChIP) experiments, we utilized BRPF1 with double-HA tag, whose cloning is explained above (see rescue experiment) in detail, was utilized. T1-160 cells were infected with lentiviral construct containing BRPF1-HA to and selected by puromycin. For ChIP, cell cross-linking was initiated by treating with formaldehyde (37%), and the reaction was halted using glycine (2.5 M). The resultant pellet, obtained after centrifugation and washing with cold PBS, was suspended in ChIP Lysis Buffer (50 mM HEPES, 1 mM EDTA, 140 mM NaCl, 1% Triton X-100, 0.1% Na-deoxycholate, 1% SDS; 1 mM PMSF; Protease Inhibitor Cocktail) and incubated on ice. Following sonication using a Bioruptor Sonicator (power setting high), the samples were centrifuged, and the supernatant containing sheared chromatin was collected. A portion of the sheared chromatin underwent decross-linking, and DNA isolation followed the ChIP clean-up kit (Zymo Research) protocols before agarose gel electrophoresis, yielding DNA fragments of 100-500 bp. Magnetic beads were washed with PBS-T, and the magnetic bead suspension was separated using a magnetic rack. The chromatin preparation underwent pre-clearing with magnetic beads, and subsequent incubation involved anti-HA-Tag Antibody (Biolegend, 901513, US) and a non-specific IgG antibody with magnetic beads. The magnetic bead-antibody complex was then incubated with pre-cleared chromatin overnight. Washing steps with low-salt (0.1% SDS,1% Triton-X 100, 2 mM EDTA, 20 mM Tris-HCl, 140 mM NaCl, pH 8.1), high-salt (0.1% SDS, 1% Triton-X 100, 2 mM EDTA, 20 mM Tris-HCl, 500 mM NaCl, pH 8.1), LiCl-containing buffers (0.25 M LiCl, 1% IGEPAL-CA 630, 1% deoxycholic acid, 1 mM EDTA, 10 mM Tris, pH 8.1), and Tris-EDTA solution ensued, with elution buffer (1% SDS and 0.1M NaHCO3) added to the magnetic beads. DNA was separated using a magnetic rack, and ChIP samples, after incubation with Rnase A, NaCl (5 M), and proteinase K, were isolated following ChIP clean-up kit (Zymo Research) protocols. Primers were designed for BRPF1-bound chromatin regions (75-100 bp PCR product) through UCSC Genome Browser and Ensembl Genome Browser and listed in **Supplementary Table 5.** Quantitative PCR (qPCR) with 2× SYBR green dye (Roche) on a LightCycler 480 (Roche) was employed for simultaneous analysis, and ChIP-qPCR results were calculated using the % input method. The initial input ratio was set at 50%, resulting in a dilution factor of 2, and the Ct value of the input DNA was adjusted accordingly.

### Analysis of Breast Cancer Patient Data

TCGA-PanCancer Atlas [37] and METABRIC datasets [38, 39] were accessed through cBioPortal [40]. Statistical analysis using One-way ANOVA was performed with Prism 8 (GraphPad Software, Inc.).

### Statistical Analysis

Analysis of EPIKOL data was performed by using the RRA method in MAGeCK. Unless otherwise stated, P values were determined by two-tailed Student’s *t*-test for all experiments in Prism 8 (GraphPad Software, Inc.), \**P*<0.05, \*\**P*<0.01, \*\*\**P*<0.001.

### DATA availability

EPIKOL screen and RNA sequencing data are deposited to the NCBI GEO database with the accession numbers GSE262577 and GSE262353.

## RESULTS

### Taxol-resistant TNBC cells demonstrate the characteristics of chemotherapy resistance

To mimic chemotherapy resistance *in vitro*, we generated Taxol (paclitaxel) resistant derivatives from SUM159PT TNBC cells. First, we measured the viability of SUM159PT cells in the presence of increasing doses of Taxol and determined IC_10_ and IC_50_ values. Then, to generate Taxol-resistant cells, SUM159PT cells were treated with IC_10_ and IC_50_ doses of Taxol for 3 days (**Fig. 1A**). Depending on the viability of the cells, they were either kept in drug-free fresh media or treated with the same dose of Taxol until they were confluent. Once the cells lost sensitivity to the drug, the amount of Taxol was doubled. This cycle was repeated until the treated cells had significantly higher IC_50_ values when compared to the starting cell population. Two different Taxol-resistant SUM159PT cells were generated by this method and named according to the starting and final doses of Taxol that they were treated with (**Fig. 1A**). T1-160 and T2-450 cells were both highly resistant to Taxol as evidenced by the increase in their IC_50_ values compared to the parental cells (**Fig. 1B**). Long-term colony formation assay in the presence of Taxol clearly demonstrated that the T1-160 and T2-450 cells can survive under high dose of Taxol treatment while the parental cells cannot (**Fig. 1C-D**). Resistant cells grew significantly slower than the parental cells (**Fig. 1E**). Immunofluorescence staining of α-tubulin on parental and resistant cells demonstrated that resistant cells were larger in size when compared to parental cells. While the morphology and microtubule organization of the parental cells markedly changed upon Taxol treatment, resistant lines were not affected, as evidenced by bright-field microscopy and α-tubulin immunofluorescence (**Fig. 1F-G**, **Supp. Fig. 1A**). Competition assay of parental and resistant cells similarly showed that only resistant cells were able to grow in the presence of Taxol (**Supp. Fig. 1B**). Annexin V/Dead cell assay demonstrated that the number of apoptotic cells was significantly higher in parental cells with Taxol treatment but there was minimal effect in resistant cells (**Fig. 1H**). Further, cleaved-PARP and increased cleaved-caspase-3 were only observed in Taxol-treated parental cells but not in resistant cells (**Fig. 1I**). Since the mechanism of action of Taxol is to stabilize microtubules, leading to cell cycle arrest at G_2_/M phase, we next assessed the cell cycle distribution patterns of Taxol-treated cells. In line with previous results, Taxol-treated parental cells significantly accumulated in the G_2_/M phase, and no major change was observed in resistant cells upon Taxol treatment (**Fig. 1J**). T1-160 and T2-450 cells were also cross-resistant to Doxorubicin and Vincristine. While Vincristine has a similar mode of action to Taxol, Doxorubicin has a completely different mechanism in which it inhibits TOP2B leading to DNA damage (**Supp. Fig. 1C**).

**Figure 1.**
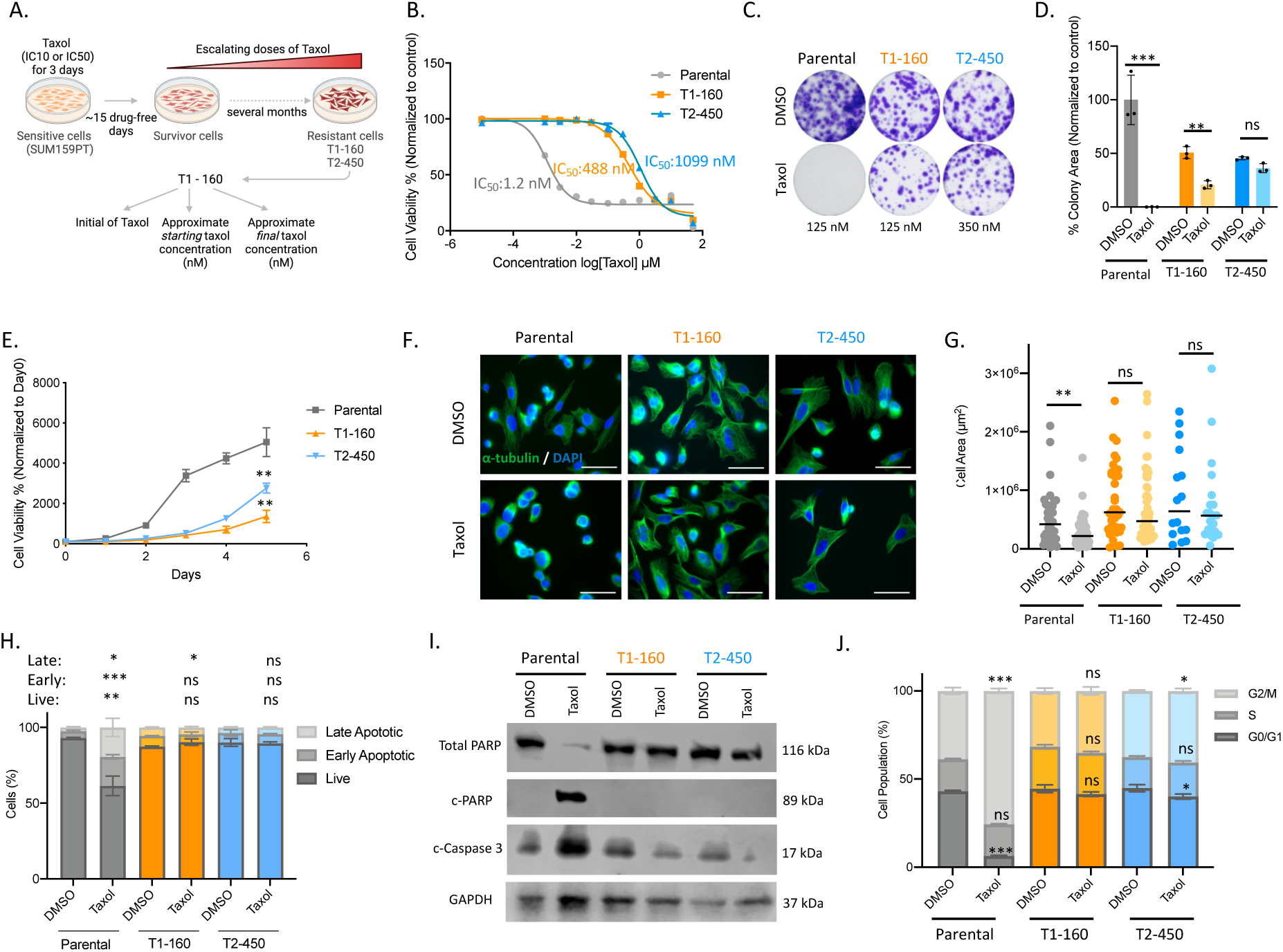
Establishment and characterization of Taxol resistant TNBC cells. **A.** Schematic representation of the protocol used for generating Taxol-resistant SUM159PT cells. The cells were exposed to increasing concentrations of Taxol (IC_10_ or IC_50_) for 72 hours, with subsequent doubling of the drug amount upon confluence. This cycle was repeated until the IC_50_ values of the cells significantly differed from the initial cell population. Figure was created with BioRender.com **B.** IC_50_ values for T1-160 (generated as a resistant cell to the folds of IC_10_ value of Taxol) and T2-450 (resistant to the folds of IC_50_). **C.** Colony formation assay in the presence of indicated amounts of Taxol **D.** Quantification of colony areas in (C.) **E.** Comparison of growth rates between parental and Taxol-resistant SUM159PT cells. **F.** Immunofluorescence staining for α-tubulin (green) and DAPI (blue) after 4 hours of exposure to DMSO or Taxol (Parental: 160 nM, T1-160: 160 nM, T2-450: 450 nM). Scale bar: 50 µm **G.** Quantification of cell area in (F.) **H.** AnnexinV/Dead cell assay after 24 hours of DMSO or Taxol treatment (Parental: 160 nM, T1-160: 160 nM, T2-450: 450 nM) **I.** Western Blot analysis of the cells shown in (H.) for Total PARP, cleaved-PARP (c-PARP), cleaved-caspase3 (c-Caspase 3) and GAPDH as a loading control. **J.** Cell cycle assay after 8 hours of DMSO or Taxol treatment (Parental: 160 nM, T1-160: 160 nM, T2-450: 450 nM). For statistical analysis, each Taxol group was compared to same cell’s DMSO group. P values determined by two-tailed Student’s t-test in comparison to control group; *p < 0.05, **p < 0.01, ***p < 0.001.

Altogether, these data showed that we successfully generated Taxol-resistant SUM159PT cells by the dose escalation method, and these cells demonstrate the characteristics of chemotherapy-resistant cells.

### Transcriptomic changes of Taxol-resistant TNBC cells highlight ABCB1-mediated resistance

To elucidate the underlying mechanisms of resistance, transcriptomes of Taxol-resistant TNBC cell lines were analyzed by RNA-sequencing. PCA plot demonstrated that triplicates of each cell line formed distinct clusters, separate from other parental or resistant cell lines (**Fig. 2A**). Comparison of T1-160 cells to the parental cells revealed 397 upregulated and 437 downregulated genes with log_2_-fold change (LFC) >2 and a significance level of p<0.001 (**Fig. 2B**). In T2-450 cells, 496 upregulated and 163 downregulated genes were identified (**Fig. 2B**).

**Figure 2.**
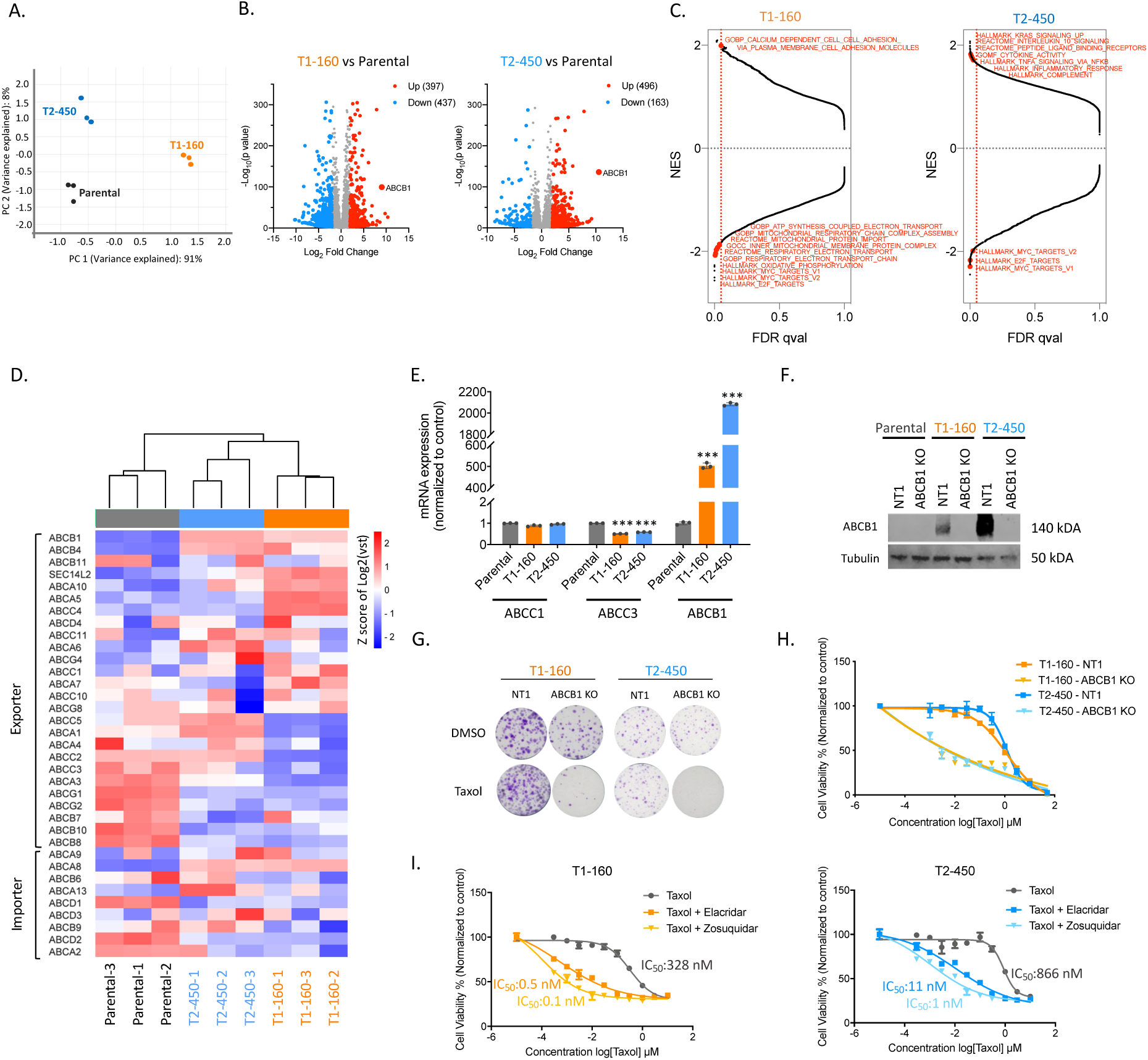
RNA-sequencing results of SUM159PT Taxol resistant cells and characterization of ABCB1-upregulated phenotype. **A.** Principal Component Analysis (PCA) plot showing the distribution of parental and Taxol resistant SUM159PT cells after RNA sequencing. **B.** Volcano plots displaying differentially expressed genes in T1-160 (left) and T2-450 (right) cells compared to parental cells, with an LFC>2 and p<0.001 cutoff. **C.** GSEA on pre-ranked gene lists based on Log_2_FoldChange values for T1-160 (left) and T2-450 (right) cells. **D.** Heatmap illustrating the gene expression pattern of multidrug transporter family members in parental and resistant cells. **E.** qPCR validations of mRNA expressions for various multidrug transporter genes. **F.** Western Blot analysis of ABCB1 protein expression in parental and resistant cells with and without ABCB1-targeting sgRNA **G.** Clonogenic assay conducted on ABCB1 knockout resistant cells in the presence of Taxol. **H.** Cell viability assay performed on ABCB1 knockout resistant cells in the presence of Taxol **I.** Cell viabilities of resistant cells upon elacridar and zosuquidar treatment with increasing doses of Taxol. P values determined by two-tailed Student’s t-test in comparison to control group; ***p < 0.001.

Gene set enrichment analysis (GSEA) in T1-160 cells revealed a predominance of negatively enriched gene sets (FDR qval < 0.05), particularly those associated with oxidative phosphorylation and the electron transport chain, indicating a potential shift towards increased glycolysis compared to parental cells. Positively enriched gene sets in T1-160 cells were linked to cell adhesion. In contrast, T2-450 cells exhibited more positively enriched gene sets, with a focus on inflammatory response and the complement system (**Fig. 2C**).

Analyzing the most significantly upregulated and downregulated genes (**Fig. 2B**), we identified ABCB1 as the top upregulated gene in both resistant cell lines. As a member of the ABC transporter family, ABCB1 is widely recognized for its strong association with drug resistance [41]. This finding aligns with previous cross-resistance results observed in the resistant cells (**Supp. Fig. 1C**), reinforcing the presence of an MDR phenotype. To provide a comprehensive overview of the ABC transporter landscape, we generated a heatmap illustrating the expression patterns of all ABC transporters, categorized as exporters and importers (**Fig. 2D**). In both resistant cell lines, the majority of exporters exhibited upregulation, while several were downregulated. Concurrently, a significant portion of importers showed downregulation, suggesting an adaptive mechanism aimed at preventing the entry of chemotherapeutic agents into the cells. Notably, ABCB1 demonstrated a significant overexpression at the RNA and protein levels (**Fig. 2E and 2F**). Given that the copy number variations (CNV) of ABCB1 are commonly linked to its elevated expression, we assessed the ABCB1 copy number in the genomic DNAs of resistant cells and identified an increase in ABCB1 copy number in both cell lines [42] (**Supp. Fig. 2A**). Knocking out ABCB1 (ABCB1 KO) in resistant cell lines completely eradicated ABCB1 expression (**Fig. 2F**). Clonogenic assays conducted with ABCB1 KO cells clearly demonstrated their inability to form colonies under Taxol pressure (**Fig. 2G**). As expected, IC_50_ values for Taxol were significantly lower in ABCB1 KO cells than controls (**Fig. 2H**). Moreover, we treated the resistant cells with three different ABCB1 inhibitors. Treatment with verapamil, a first-generation ABC transporter inhibitor [6] and calcium channel blocker, significantly lowered IC_50_ values for Taxol (**Supp. Fig. 2B**). Fluorescence imaging of Calcein-AM further illustrated that the uptake of Calcein was restricted in Taxol-resistant cells but augmented in the presence of verapamil, indicating the functional role of ABCB1 in these resistant cells (**Supp. Fig. 2C**). Elacridar, a second-generation ABC inhibitor that selectively binds to several ABC proteins, and zosuquidar, a third-generation ABC inhibitor with high affinity for ABCB1 [43], both reversed resistance, with the most pronounced effect observed with zosuquidar (**Fig. 2I**). This set of data underscores the significant impact of ABCB1 on the resistance of these cells.

### Chromatin-focused screens reveal epigenetic vulnerabilities of resistant cells

To uncover epigenetic regulators of Taxol resistance, we utilized two complementary approaches: epigenome-wide CRISPR/Cas9 and epigenetic probe-library screens. For the CRISPR/Cas9 library screen, we took advantage of our previously published chromatin-focused sgRNA library (EPIKOL) [30]. Following infection and puromycin selection, T1-160 cells were subdivided into DMSO and Taxol-treated groups. Subsequent to culturing period and next-generation sequencing (NGS), the Taxol-treated samples were compared to the DMSO-treated group to identify epigenetic modifiers whose loss revert resistance (**Fig. 3A**). ABCB1 and ABCG2, positive controls present in our library, were highly depleted in the presence of Taxol, validating the reliability of our screen. Notably, members of COMPASS/MLL complex, SWI/SNF complex, and de-ubiquitination related genes were among the most significantly depleted genes (**Fig. 3B**).

**Figure 3.**
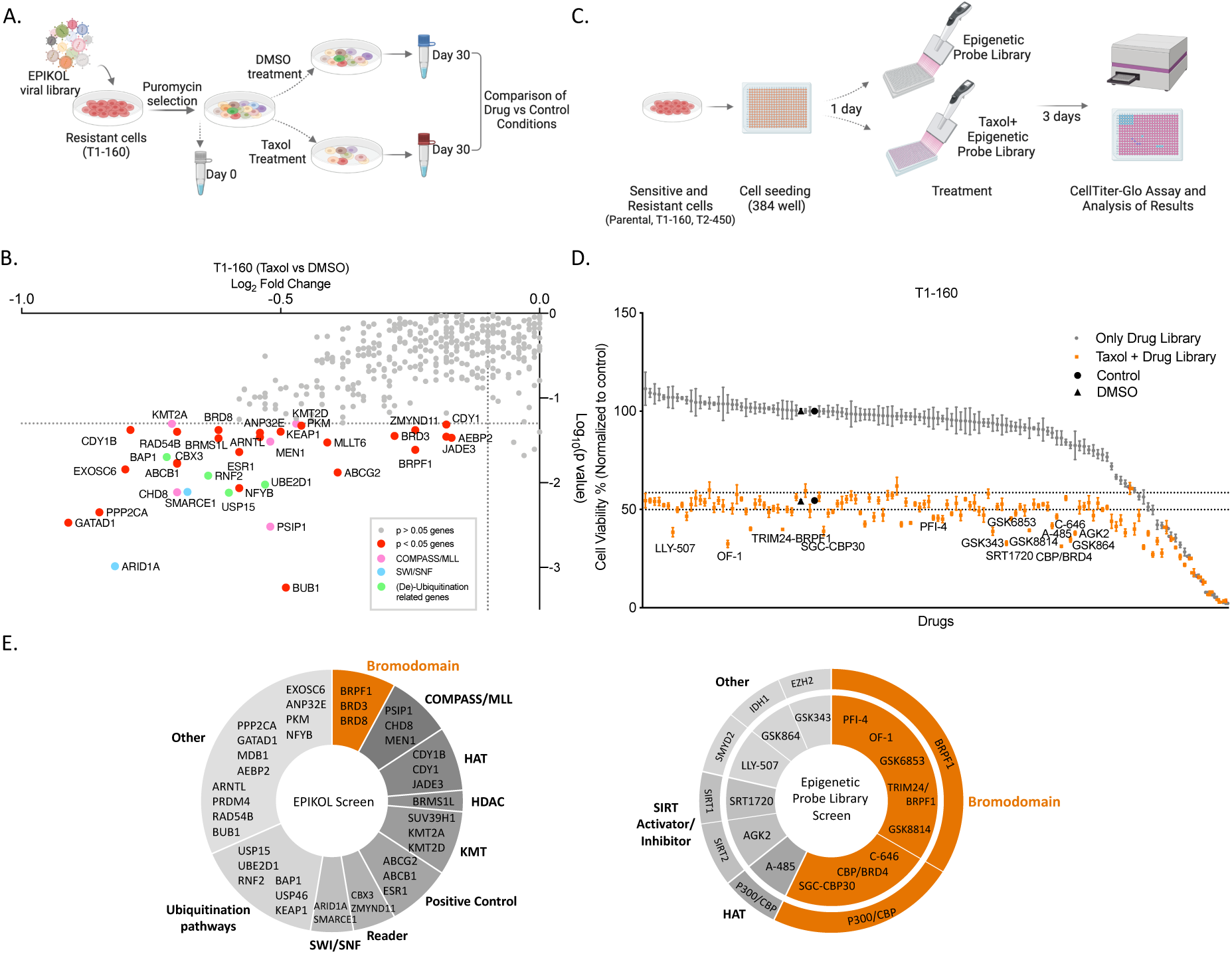
Chromatin-focused genetic and chemical probe library screens reveal epigenetic vulnerabilities of Taxol resistant TNBC cells. **A.** Schematic representation of EPIKOL screen on T1-160 resistant cells. Cells were infected with 1000x representation of library with MOI=0.3 and selected with puromycin. Following completion of Cas9 activity approximately 9 days after transduction, cells were divided into two groups as DMSO-treated and Taxol-treated (160 nM) groups. The final time-point samples were compared to identify epigenetic modifiers that sensitize Taxol-resistant cells. Figure was created with BioRender.com **B.** Result of the EPIKOL screen performed on T1-160 cells, presented as Log_2_FoldChanges of genes in Taxol-treated samples compared to the DMSO-treated group. Genes that have p<0.05 were colored and labeled **C.** Schematic representation of epigenetic chemical probe library screen. Cells were seeded to 384-well plates as 750 cells/well and next day treated with Taxol (Parental: 1.5 nM, T1-160: 300 nM, T2-450: 500 nM) and 1x concentration of epigenetic probe library. After 3 days, cell viability was measured. Figure was created with BioRender.com. **D.** Results of epigenetic probe library screen performed on T1-160 cells. Grey dots represent the effect of epigenetic probes alone while orange dots show changes in cell viability when Taxol was combined with a specific epigenetic probe. Epigenetic probes that significantly reversed drug resistance are labeled on the graph. If the negative control of a given probe also showed a significant depletion in cell viability, neither of the probes was labeled on the graph. **E.** Classification of hits identified in genetic (left) and chemical screens (right). For chemical screen, target gene of chemical probes were indicated in the outer circle.

As an alternative approach, we utilized an epigenetic-probe library consisting 117 chemical probes targeting a broad range of epigenetic factors (**Supp. Fig. 3A**). Parental and resistant cells were treated with Taxol alone or in combination with epigenetic probes. After 72 hours of treatment, cell viability was assessed (**Fig. 3C**). The majority of the epigenetic probes exhibited minimal impact on T1-160 cell viability individually (**Fig. 3D**). Various bromodomain inhibitors, SIRT activator/inhibitors, mIDH1 inhibitor, as well as EZH2 inhibitor significantly decreased cell viability when applied together with Taxol (**Fig 3D**).

Additionally, we performed the probe-library screen on SUM159PT parental and T2-450 Taxol-resistant cells **(Supp. Fig. 3B**-C). Similar to the T1-160 cells, numerous bromodomain inhibitors, particularly those targeting BRPF1, and SIRT activator/inhibitors affected cell viability when combined with Taxol. Interestingly, members of the lysine methyltransferase family exhibited sensitizing effects specifically in T2-450 cells **(Supp. Fig. 3D)**. The majority of the epigenetic-probes that reduced cell viability in combination with Taxol were shared between T1-160 and T2-450 cells **(Supp. Fig. 3E)**. Notably, inhibitors such as PFI-4, OF-1, GSK6853, and TRIM24/BRPF1 demonstrated a substantial impact on cell viability when combined with Taxol in resistant cells. In contrast, these BRPF1 inhibitors did not have a significant effect on parental cells.

To prioritize the hits from genetic and chemical screens, we classified the genes and inhibitors according to their complexes or epigenetic modifier classes (**Fig. 3E**). EPIKOL screen hit genes (p<0.05) were classified into nine main categories: bromodomain-containing proteins, COMPASS/MLL complex, Histone acetyltransferase (HAT), Histone deacetylase (HDAC), Lysine methyltransferase (KMT), readers, SWI/SNF complex, ubiquitination pathways, and others. Epigenetic-probe library screen hit targets include BRPF1 and p300/CBP of bromodomain family proteins, SIRT1, SIRT2, SMYD2, IDH1, EZH2. Notably, when both of the screen results are considered together, BRPF1, a member of the bromodomain family, emerged as the sole gene identified as a hit in both screens.

### EPIKOL screen hits are validated through functional assays

Genes that have high log-fold changes and that are not previously associated with chemotherapy resistance were selected for further validation (**Fig. 3B and Supp. Fig. 4A**). BRPF1 was included in this group as it was the only gene that was identified in both screens. To validate EPIKOL screen hits, we performed a dual-color cell growth competition assay (**Fig. 4A**). For this, mCherry labeled T1-160-Cas9 stable cells were infected with NT1 and mixed with eGFP labeled T1-160 Cas9 stable cells carrying sgRNA of interest in a 1:1 ratio. For 24 days, cells carrying sgRNAs against the selected hits were outcompeted by the cells carrying non-targeting sgRNA (NT1). ABCB1 KO served as a positive control, exhibiting the most pronounced effect in the presence of Taxol (**Fig. 4B-C**). In this assay, knockouts of GATAD1, PPP2CA, CHD8, BRD8, SMARCE1, KMT2A, and MEN1 resulted in significant decreases in cell viability (**Supp**. Fig. 4B). Despite BRPF1 not being the top scorer in the EPIKOL screen, it consistently exerted a small yet significant effect on cell viability in combination with Taxol. This phenotype was validated with three independent sgRNAs targeting BRPF1 (**Fig. 4B-C**). Given that BRPF1 was the only gene demonstrating a phenotypic effect through both genetic (knockout) and pharmacological (inhibition with small compounds) approaches, we chose to further focus on BRPF1 as a potential regulator of chemoresistance.

**Figure 4.**
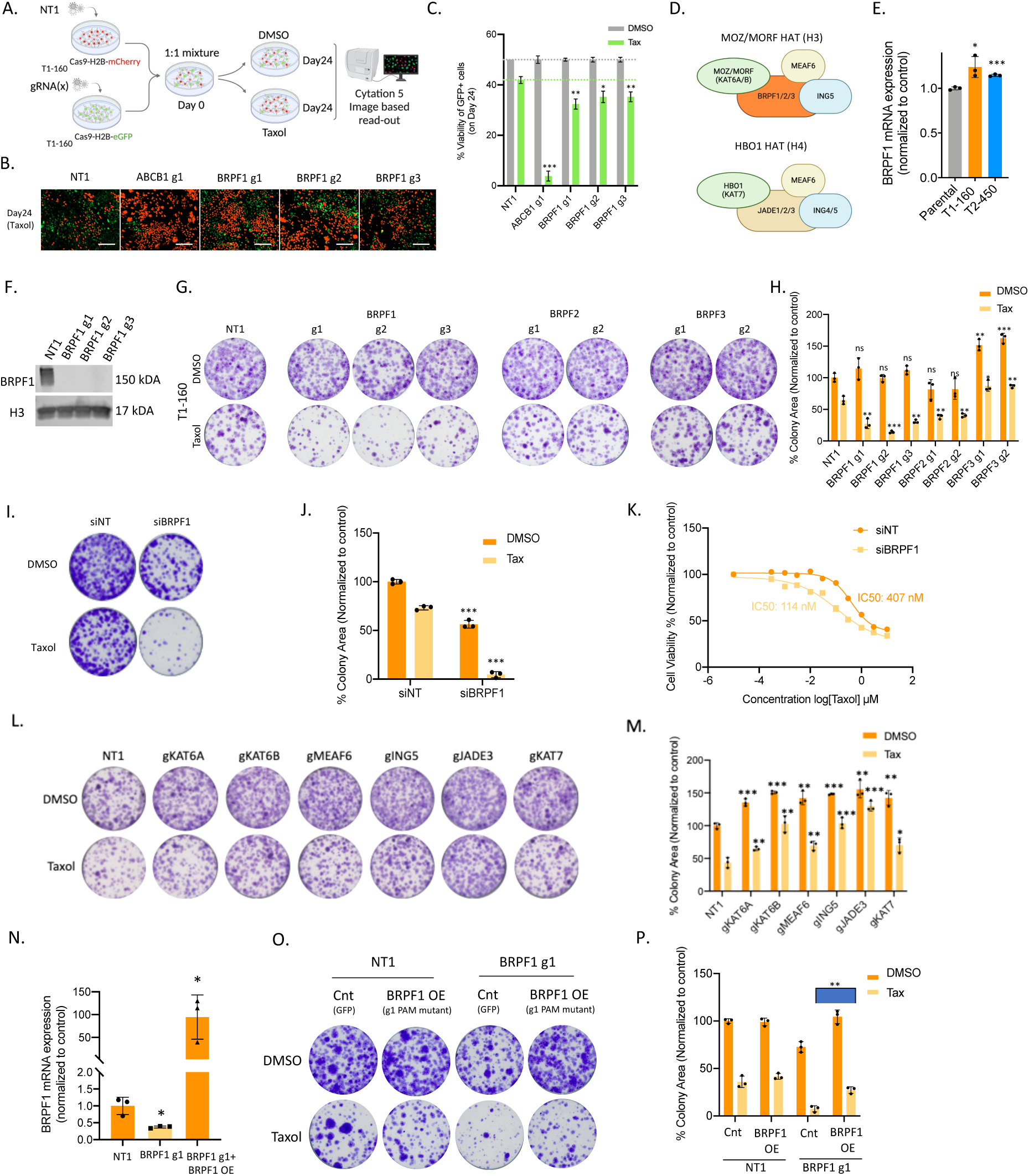
Genetic perturbation of BRPF1 sensitizes Taxol-resistant T1-160 cells. **A.** Schematic representation of dual-color competition assay. mCherry-H2B or eGFP-H2B labeled T1-160 cells were infected with NT1 or sgRNA of interest. Cells were mixed in a 1:1 ratio, treated with Taxol (160 nM) and imaged after attachment and every 4 days until Day 24. Ratio of eGFP-H2B-T1-160 cells carrying sgRNA of interest were determined to identify the genes whose loss revert resistance. Figure was created with BioRender.com **B.** Representative images of dual-color competition assay for knockouts of indicated genes. Scale bar=200 µm **C.** Statistical analysis of eGFP+ cells upon ABCB1 and BRPF1 knockout after dual-color competition assay **D.** Complexes that are formed by BRPF and JADE proteins. **E.** BRPF1 expression levels in parental and resistant cells. Figure was created with BioRender.com **F.** Western blot analysis of BRPF1 expression upon BRPF1 knockout in T1-160 cells. **G.** Clonogenic assay showing the effect of BRPF1, BRPF2, and BRPF3 knockouts in the presence of Taxol in T1-160 cells. **H.** Quantification of the colonies in (G.). **I.** Effect of BRPF1 siRNA on colony forming abilities of T1-160 cells in the presence of Taxol. **J.** Quantification of colonies in (I.). **K.** Cell viability measurement performed with siNT and siBRPF1 samples in the presence of Taxol in T1-160 cells. **L.** Effect of MOZ/MORF complex members’ knockouts on colony forming abilities of T1-160 cells in the presence of Taxol. **M.** Quantification of colonies in (L.). **N**. BRPF1 mRNA levels upon knockout and overexpression of PAM mutant version of BRPF1 in T1-160 cells **O.** Rescue experiment with overexpression of PAM mutant version of BRPF1 on BRPF1 knockout cells in the presence of Taxol. **P.** Quantification of colonies in (O.). P values determined by two-tailed Student’s t-test in comparison to control group; *p < 0.05, **p < 0.01, ***p < 0.001.

### Knockout of BRPF1 sensitizes Taxol-resistant cells

Bromodomain and PHD Finger Containing 1 (BRPF1, also known as peregrin and BR140) is a member of the Histone acetyltransferase (HAT) complex containing Monocytic leukemia zinc finger protein (MOZ, KAT6A) or monocytic leukemia zinc finger protein-related factor (MORF, KAT6B) as catalytic subunits **(Fig. 4D**) [44]. BRPF1 is a chromatin reader that contains a plant homeodomain (PHD)-zinc-knuckle-PHD (PZP) module at the N-terminus, followed by a bromodomain recognizing acetyl-lysines, and a C-terminal proline-tryptophan-tryptophan-proline (PWWP) domain. BRPF1 acts as a scaffold by binding to MOZ/MORF through its N-terminus motifs and ING5 and MEAF6 through the downstream motif [45, 46].

Initially, we assessed the expression levels of BRPF1 and observed a slight upregulation in BRPF1 mRNA levels in resistant cells (**Fig. 4E**). We confirmed knockout efficiency of the BRPF1-targeting sgRNAs used through qPCR and western blotting. Despite modest decreases in mRNA levels with sgRNA expression, BRPF1 protein was completely lost in KO samples when compared to non-targeting sgRNA controls (**Supp. Fig. 5A**, **Fig. 4F**). Notably, BRPF1 has two structurally close proteins, BRPF2 (BRD1) and BRPF3, which are known to be part of the MOZ/MORF complex [47]. Although no depletion was observed in BRPF2 and BRPF3 sgRNAs during the EPIKOL screen, to exclude the possibility of their involvement in Taxol resistance, we also performed knockouts for BRPF2 and BRPF3 and analyzed their effects on Taxol resistance (**Supp. Fig. 5B**). As evidenced by the clonogenic assay, the most significant effect in the presence of Taxol was observed in BRPF1 KO samples, with BRPF2 and BRPF3 KOs having minor or no effects (**Fig. 4G-H**). In the second resistant line, T2-450, loss of BRPF2 and BRPF3 did not show any effects, whereas BRPF1 was essential for cell survival (**Supp. Fig. 5C**-D). These findings suggest that the sensitization effect to Taxol is primarily attributed to the knockout of BRPF1.

As an independent loss of function approach, we utilized siRNA targeting BRPF1, which resulted in a substantial reduction in BRPF1 mRNA levels within a short timeframe (24-48 hours) (**Supp. Fig. 5E**). This rapid downregulation mirrored the sensitization phenotype observed in knockout samples in the presence of Taxol (**Fig. 4I-J**). The acute suppression of BRPF1 enabled us to assess sensitization in siBRPF1 samples through an ATP-based cell viability assay, clearly demonstrating the nearly 4-fold change in IC_50_ values (**Fig. 4K**). Importantly, BRPF1 knockdown in parental cells did not significantly alter the IC_50_ value (**Supp. Fig. 5F**-G).

To investigate whether the observed phenotype is solely dependent on BRPF1 or extends to the entire MOZ/MORF complex, we performed knockouts for all complex members (**Supp. Fig. 5H**). We included the members of HBO1 complex since the two complexes have shared members. In contrast to BRPF1, none of the complex members exhibited sensitization upon knockout in the presence of Taxol (**Fig. 4L-M**). To further confirm the role of BRPF1 on Taxol resistance, we conducted a rescue experiment by reintroducing a PAM mutant version of BRPF1 into the knockout background (**Fig. 4N**). The ability to form colonies in the presence of Taxol was restored when BRPF1 was re-expressed after the knockout (**Fig. 4O-P**). Overall, these findings suggest that BRPF1 has a significant role in Taxol resistance of TNBC cells.

To understand the clinical significance of BRPF1 alongside the members of MOZ/MORF and HBO1 complexes, we analyzed publicly available breast cancer patient data from TCGA which includes all breast cancer subtypes such as Luminal A, Luminal B, Her2 and Basal subtype. Our findings indicated that, in the Basal subtype containing TNBC, there is a distinct upregulation solely in BRPF1 mRNA levels, in contrast to other members of the MOZ/MORF and HBO1 complexes. (**Supp. Fig. 6A**). Moreover, we observed a positive correlation between BRPF1 expression and increasing breast cancer grade, distinguishing BRPF1 from other members (**Supp. Fig. 6B**). These results indicate that BRPF1 could potentially function as a prognostic marker in breast cancer patients.

### BRPF1 inhibitors recapitulate the effects of BRPF loss on Taxol-resistant cells

Through the unbiased chemical probe library screen (**Fig. 3D**), several BRPF1 inhibitors demonstrated the potential to sensitize Taxol-resistant cells. Among them, PFI-4 (targets BRPF1b), and OF-1 (targets pan-BRPF) are structurally different (**Fig. 5A**). These inhibitors significantly reduced cell viability when combined with Taxol in resistant cells, with no impact on parental cells (**Fig. 5B**). GSK6853 was another BRPF1-specific inhibitor identified as a potential hit, however it had a modest effect on cell viability of T1-160 with no effect on parental and T2-450 cells (**Supp. Fig. 7A**). We also tested the effects of GSK5959, which has a chemical structure similar to the PFI-4’s but was not present in the library. GSK5959 sensitized T1-160 cells to Taxol, and its effect on T2-450 cells was similar to the results observed on knockout samples decreasing cell viability even in the absence of Taxol (**Supp. Fig. 7A**). For subsequent experiments, we mainly utilized PFI-4 and OF-1 as they demonstrated consistent efficacy in both cell lines and were identified during the initial library screen. Clonogenic assay on T1-160 and T2-450 cells demonstrated that combination of Taxol with PFI-4 and OF-1 impaired cell viability of resistant cells with minimal impact on parental cells (**Fig. 5C-D**). Bright field images of Taxol and PFI-4 and OF-1 treated cells also illustrated the abnormal cell shapes and lower number of cells when compared to controls (**Supp. Fig. 7B**). Increased cleavage of PARP was observed in Taxol-treated parental cells and combination-treated T1-160 and T2-450 cells (**Fig. 5E**). Immunofluorescence staining of α-tubulin on T1-160 cells in the presence of Taxol and BRPF1 inhibitors demonstrated round cells with shrunk microtubules (**Fig. 5F-G**).

**Figure 5:**
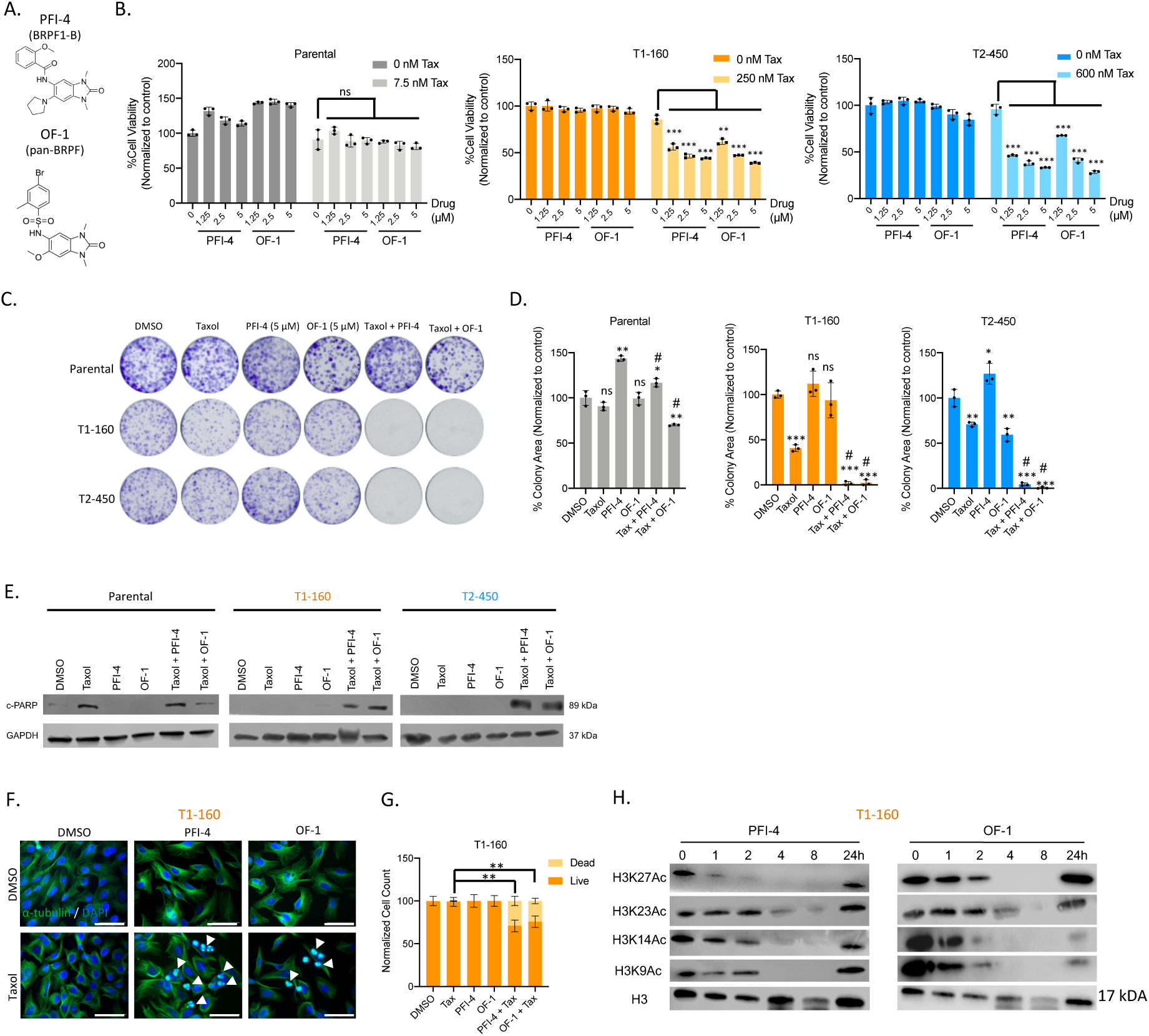
Chemical probe library screens reveal BRPF1 inhibitors as Taxol sensitizers. **A.** Chemical structures of BRPF1 inhibitors identified during probe library screen. PFI-4 specifically targets BRPF-1 and OF-1 is a pan-BRPF inhibitor acting against BRPF1/BRPF2/BRPF3. **B.** Validations of the effect of PFI-4 and OF-1 in combination with Taxol on parental, T1-160 and T2-450 cells. **C.** Clonogenic assay results of Taxol (3 nM for Parental, 125 nM for T1-160 and 300 nM for T2-450) and BRPF1i (5 µM each) combinations **D.** Quantification of colonies in (C.). P values determined by two-tailed Student’s t-test in comparison to DMSO; *p < 0.05, **p < 0.01, ***p < 0.001. # indicates p<0.01 and # indicates p<0.001 in comparison to Taxol. **E.** Western Blot analysis for cleaved-PARP (c-PARP) and GAPDH as loading control upon combination treatment. **F.** Immunofluorescence staining with α-tubulin (green) and DAPI (blue) in the presence of Taxol (Parental: 160 nM, T1-160: 160 nM, T2-450: 450 nM) and/or BRPF1 inhibitors (5 µM each) for 16 hours. **G.** Quantification of dead cells in (F.). Scale bar: 50 µm **H.** Western blot analysis of PFI-4 and OF-1 treated T1-160 cells on different timepoints for histone acetyl marks.

Lastly, we checked if PFI-4 or OF-1 impairs the functioning of the BRPF1 complex. BRPF1 regulates the acetylation of H3K9, H3K14, and H3K23 by increasing the HAT activity of the catalytic subunits MOZ and MORF upon assembly of the complex. Through its bromodomain, BRPF1 also reads H2AK5Ac, H4K12Ac, and H3K14Ac [48]. Treatment of PFI-4 and OF-1 decreased H3K14Ac and H3K9Ac levels as early as 4 hours, while loss of H3K23Ac took around 8 hours (**Fig. 5H**). Open chromatin marker H3K27Ac also decreased upon PFI-4 and OF-1 treatment. Almost all of the acetylation marks were restored within 24 hours.

Although knockout of MOZ/MORF complex members did not affect cell viability, we evaluated KAT6A and KAT6B inhibitors for potential effects on Taxol resistance (**Supp. Fig. 7C**). While KAT6A inhibition alone with WM-1119 had no significant effect on cell viability when combined with Taxol, WM-8014 (KAT6A and KAT6B inhibitor) showed a small decrease. However, the efficacy of KAT6A/B inhibitors did not match that of PFI-4 and OF-1, suggesting that the reader function of BRPF1 may be more crucial than its scaffolding function for the HAT activity of the complex. In summary, our findings indicate that BRPF1 inhibitors, when combined with Taxol, significantly decrease cell viability in resistant cells.

### Transcriptomic changes caused by BRPF1 reveal defects in translation machinery in Taxol-resistant cells

To gain insight into the transcriptomic changes caused by BRPF1, we treated T1-160 cells with PFI-4 and OF-1 or knocked out BRPF1 and performed RNA-sequencing. PFI-4 caused upregulation of 327 genes and downregulation of 336 genes, while OF-1 had a greater impact on transcriptome with over 2000 genes being either upregulated or downregulated (**Fig. 6A**). We then performed overlap analysis with biological processes from the Molecular Signature Database (MsigDB) on commonly downregulated or upregulated genes upon inhibitor treatment (**Fig. 6B**). Programmed cell death was one of the upregulated signatures upon inhibitor treatment. Notably, commonly downregulated genes significantly overlapped with numerous RNA and ribosome biogenesis-related pathways. BRPF1 targeting sgRNA #2 resulted in 382 upregulated and 719 downregulated genes. sgRNA #3 caused upregulation and downregulation of approximately 200 genes (**Fig. 6C**). Majority of downregulated genes in BRPF1 sgRNA #3 were common with sgRNA #2, and these genes significantly overlapped with ribosome biogenesis and translation-related pathways (**Fig. 6D**). As a result, inhibition or knockout of BRPF1 significantly decreased the expression of ribosome biogenesis and translation-related genes, suggesting an impairment of translation machinery. We observed a slight but consistent decrease in the mRNA levels of nearly all ribosome-related genes upon BRPF1 knockout (**Fig. 6E and Supp. Fig. 8A**) and inhibition (**Supp. Fig. 8B**). To assess the functional role of downregulation of ribosomal genes, we measured global translation rate with sunset assay [49] (**Fig. 6F**). Notably, all BRPF1 knockout T1-160 cells had lesser amount of puromycin labeled proteins when compared to control, suggesting that the translation rate is decreased upon BRPF1 loss.

**Figure 6:**
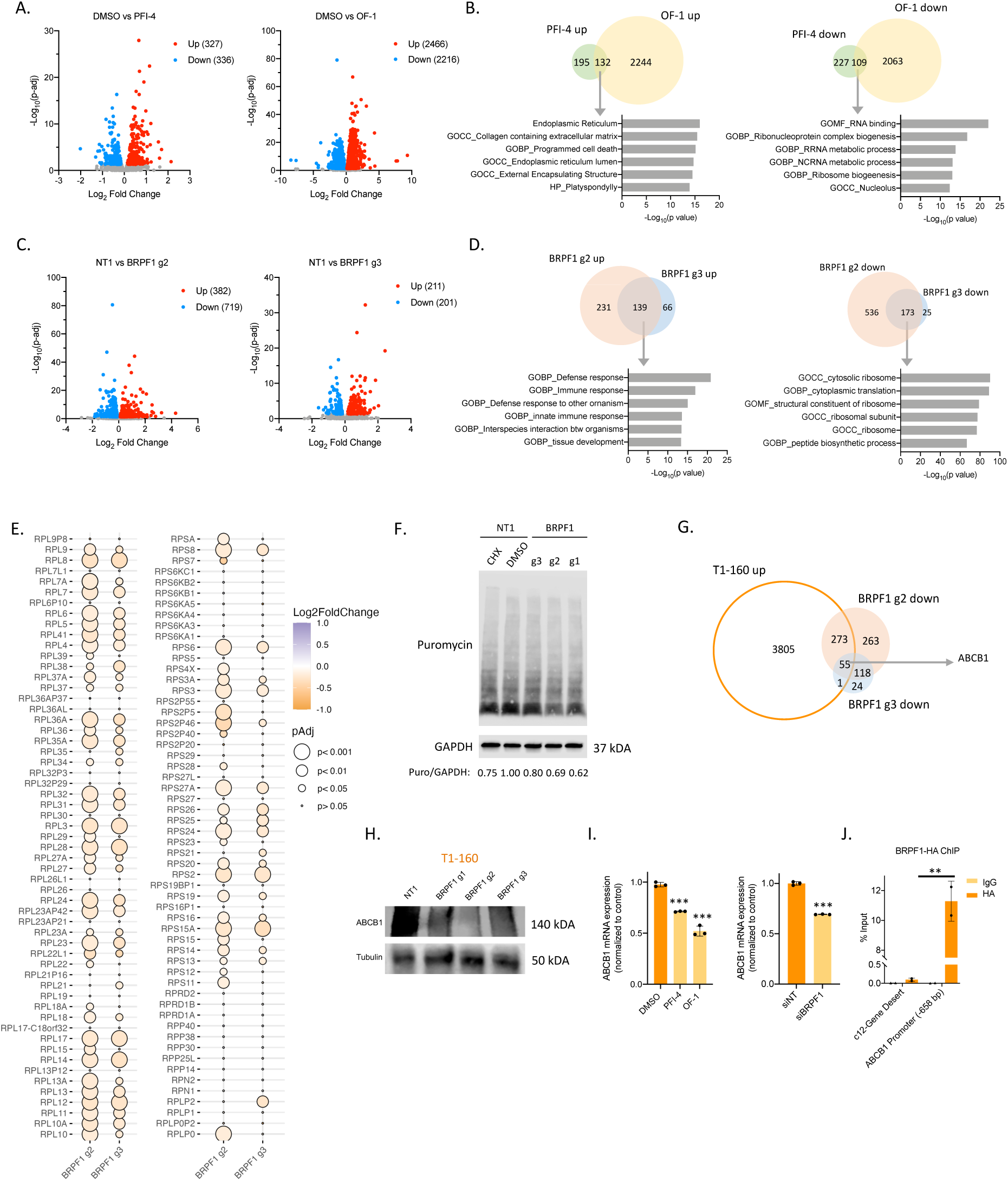
Transcriptomic changes caused by BRPF1 inhibition indicate ribosome biogenesis defects. **A.** Volcano plots showing differentially expressed genes (p<0.05) after 72 hours of PFI-4 (5 µM) (left)and OF-1 (5 µM) (right) treatment on T1-160 cells. **B.** Venn diagrams indicate the numbers of common upregulated or downregulated genes in PFI-4 or OF-1treated cells. Significant pathways of the common upregulated and downregulated genes were identified by the overlap analysis via MsigDB. **C.** Volcano plots showing differentially expressed genes (p<0.05) in BRPF1 knockout T1-160 cells on post-transduction day 12. **D.** Analysis of common upregulated and downregulated genes in BRPF1 knockout cells and overlap analysis for gene ontology. **E.** Balloon plot depicting the Log_2_FoldChanges and adjusted-p values of ribosome-related genes in BRPF1 knockout cells. **F.** SUnSET assay showing global translation levels in control and BRPF1 knockout cells. Band intensities were normalized loading control GAPDH and ratio of NT1 sample was normalized to 1. **G.** Comparison of the genes upregulated in T1-160 cells and downregulated in BRPF1 knockout T1-160 cells. **H**. Western Blot showing ABCB1 protein levels in BRPF1 Knockout T1-160 cells. **I.** ABCB1 mRNA levels upon 72 hours of PFI-4 and OF-1 and 48 hours of siBRPF1 treatment on T1-160 cells. **J.** ChIP-qPCR analysis of BRPF1 binding on ABCB1 promoter in HA-tagged BRPF1 expressing-T1-160 cells. Ch12-Gene Desert region serves as a negative control. Data is representative of 3 independent biological replicates. P values determined by two-tailed Student’s t-test in comparison to control group; *p < 0.05, **p < 0.01, ***p < 0.001.

Next, we focused on the set of genes upregulated in resistant cells, that were downregulated upon BRPF1 knockout (**Fig. 6G**). Notably, *ABCB1* was among this set of genes. ABCB1 protein was expressed at significantly lower levels in BRPF1 KO-resistant cells (**Fig. 6H**). In addition, we observed significant downregulation of the ABCB1 expression in Taxol-resistant cells treated with either BRPF1 inhibitors or siRNA (**Fig. 6I**).

We investigated whether BRPF1 directly binds to the ABCB1 promoter to facilitate its transcription and performed chromatin immunoprecipitation (ChIP). We used anti-HA antibody to pull down HA-tagged BRPF1 and demonstrated a significant enrichment of BRPF1 on the ABCB1 promoter when compared to IgG and chromosome 12 gene desert region (**Fig. 6J**) [50]. Overall, these findings provide strong support for the direct responsibility of BRPF1 in regulating ABCB1-mediated Taxol resistance.

## DISCUSSION

The present study aimed to unravel the epigenetic mechanisms underlying Taxol resistance in TNBC cells. To recapitulate chemotherapy resistance *in vitro*, two independent Taxol-resistant TNBC cell lines (T1-160 and T2-450) were generated through a dose escalation method. These resistant cells demonstrated characteristics typical of chemotherapy resistance, such as increased IC_50_ values, slower growth, altered cell morphology, and resistance to apoptosis induction compared to parental cells **(Supp. Fig. 9)**. Transcriptomic analyses revealed distinct characteristics in each cell line, such as decreased oxidative phosphorylation in T1-160 [51–53] and upregulation of inflammatory response-related pathways in T2-450 [54]. Notably, both cell lines exhibited elevated expression of ABCB1, an ATP-binding cassette transporter associated with MDR, which was further validated through functional assays emphasizing the role of ABCB1 in our Taxol-resistant TNBC cell lines.

Despite successfully reversing resistance in T1-160 cells using various ABCB1 inhibitors, such as verapamil, elacridar, and zosuquidar, the high toxicity and limited benefits of these inhibitors preclude their use in combination therapies. CNV is known to contribute to drug resistance; however, recent studies indicated that although the copy number of *ABCB1* was elevated, it was insufficient to activate *ABCB1* expression without the transcriptional regulation, emphasizing the inadequacy of CNV alone [14, 21, 29, 55]. Thus, understanding the upstream regulators of ABCB1 specific to cancer cells is crucial for developing effective and safe treatment strategies.

To explore potential epigenetic regulators contributing to Taxol resistance, comprehensive epigenome-wide CRISPR/Cas9 and epigenetic probe-library screens were employed. The convergence of results from both screens pinpointed BRPF1 as a significant hit. BRPF1 orchestrates histone acetylation by bringing the MEAF6 and ING5 accessory proteins and MOZ/MORF catalytic subunits together. It is known that BRPF1 is indispensable during embryonic development [56, 57] and causes intellectual disability when mutated [58]. In the context of cancer, upregulation of BRPF1 is associated with low survival rates in HCC patients [48]. Analysis of TCGA data revealed BRPF1 mutations or CNVs in various cancer types [44, 59]. However, the link between BRPF1 and chemoresistance was largely unexplored. BRPF2 and BRPF3, close paralogs of BRPF1, may also form complexes with MOZ/MORF. BRPF1 knockout significantly impaired colony-forming abilities in Taxol-resistant cells in the presence of Taxol, surpassing the impact of BRPF2 or BRPF3 knockouts. In T2-450 cells, BRPF1 loss impaired colony-forming ability even in the absence of Taxol, suggesting that BRPF1 was required for cell fitness of Taxol-resistant T2-450 cells. Intriguingly, the catalytic or accessory subunits of the MOZ/MORF complex showed no effect, emphasizing the unique role of BRPF1 as a reader in the chemoresistance context. Indeed, the rescue of BRPF1 knockout by overexpressing the PAM mutant version of BRPF1 was sufficient to restore Taxol resistance of T1-160 cells.

Additionally, we investigated BRPF1 inhibitors, PFI-4 and OF-1, revealing consistent efficacy in reducing cell viability when combined with Taxol, suggesting their potential as future therapeutic agents. Transcriptomic changes induced by BRPF1 loss or inhibition provided valuable insights into defects in Taxol-resistant cells, particularly the downregulation of ribosomal and translation-related genes. Functional assays further demonstrated a decreased global translation rate in BRPF1 knockout cells, linking BRPF1 to the regulation of the translation machinery in Taxol-resistant TNBC cell lines.

RUNX2, a downstream target of BRPF1, has been reported to negatively regulate the transcriptional control of rRNA genes [60, 61]. The deficiency of another BRPF1 target, RUNX1, decreases ribosome biogenesis and translation, causing hematopoietic stem and progenitor cells to develop resistance to genotoxic stress [62]. The discrepancy on the effects of RUNX transcription factors (TFs) might result from the chromatin regulatory factors with which they associate [63]. It has been reported that MOZ and MORF function as transcriptional coactivators for RUNX TFs [50]. Although BRPF1 was not previously associated with ribosome biogenesis, our findings suggest that BRPF1 has a direct role in the regulation of translation machinery in Taxol-resistant TNBC cell lines. It is possible that BRPF1, as the scaffolding member of the complex that directs it to specific genomic locations, might be regulating the interaction between MOZ/MORF and RUNX TFs in the chemoresistance context.

In this study, through genetic and chemical screens, we identified BRPF1 as a critical regulator of Taxol-resistance in TNBC cells. The depletion of BRPF1 through CRISPR/Cas9 or siRNA, as well as inhibition using PFI-4 or OF-1, resulted in a reduction of ABCB1 expression. The regulation of ABCB1 levels by BRPF1 involved its direct binding to the *ABCB1* promoter (**Fig. 7**). Analysis of publicly available breast cancer patient data also revealed elevated expression of BRPF1 in basal subtype as well as in higher grade tumors, indicating a role for BRPF1 in disease progression. Additionally, in an independent study, we demonstrated that castration-resistant prostate cancer cells also depend on BRPF1 in docetaxel and cabazitaxel resistance, highlighting BRPF1’s role as an *ABCB1* regulator controlling mTOR and UPR signaling [64]. In this present study, we unraveled that BRPF1 deficiency led to diminished ribosome biogenesis, causing a decline in the global translation rate, potentially contributing to an overall decrease in ABCB1 levels in resistant cells. The intricate interplay between BRPF1, ribosome biogenesis, and ABCB1 expression elucidates a novel mechanism underlying Taxol resistance in TNBC cells and holds promise as a therapeutic strategy, particularly in patients exhibiting elevated ABCB1 levels.

**Figure 7:**
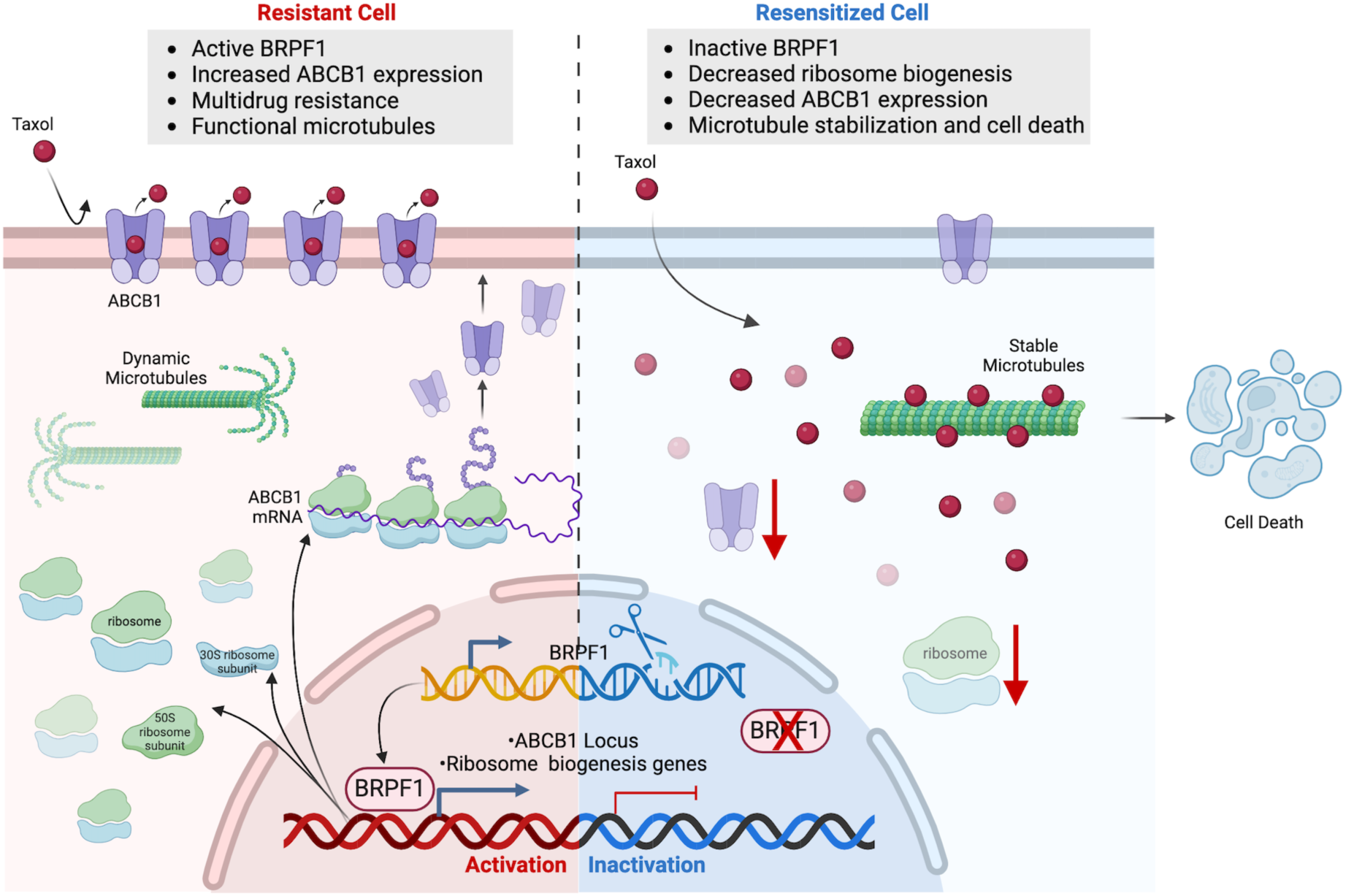
Proposed model of Taxol resistance regulation by BRPF1. The role of BRPF1 in Taxol resistance was summarized. Figure was created with BioRender.com

## Author Contributions

Study design: TB-O, OY-B, TTO; data generation: OY-B, AC, EY, NP-D, MP, BD; data analysis: OY-B, ACA, ADC, AC, HS; chemical library formation: MP, AC, UO; data interpretation: OY-B, TB-O, TTO, NAL, BC, CAA, UO; initial manuscript draft: OYB, TB-O and TTO; approved final manuscript: all authors.

## Acknowledgements.

Financial support was obtained from The Scientific and Technological Research Council of Turkey (TUBITAK) (1003-216S461 and 1001-221S419 Grants). We thank Dr. Alişan Kayabolen and Rauf Günsay for their help during plasmid design and cloning experiments. Schematic representation figures were created in BioRender.com and licensed for publication (Fig1: MB26MRYP6N, Fig3: KV26MS22YT, LG26MS1IXD, Fig4: NF26MS3F98, FL26MS2T21, Fig7: QS26MS3UDF, Supp. Fig.9: BL26MS4A1T). The authors gratefully acknowledge the use of the services and facilities of the Koç University Research Center for Translational Medicine (KUTTAM), funded by the Presidency of Turkey, Head of Strategy and Budget.

