## Supplementary methods, tables and figures for "Chromatin-focused genetic and chemical screens identify BRPF1 as a targetable vulnerability in Taxol-resistant triple-negative breast cancer"

### SUPPLEMENTARY INFORMATION

### SUPPLEMENTARY MATERIALS AND METHODS

**Live Cell Microscopy.** Cells were seeded as 75.000 cells/well in 12-well plates and next day treated with Taxol; 160 nM for Parental and T1-160 and 450 nM for T2-450. For drug combination experiments, concentrations of PFI-4 and OF-1 was 5 µM. After drug treatment, phase contrast images were taken as 2x2 with 10x objectives using Cytation5 (Biotek, USA) every 15 minutes for 72 hours.

**Fluorescent-labeled resistant cell assay.** Parental cells were transduced with PGK-H2BeGFP (Addgene #21210) viruses, resistant cells were transduced with PGK-H2BmCherry (Addgene #21217) at MOI~5 ensure each cell was fluorescently labeled. Parental and resistant cells were mixed in a 1:1 ratio as 40.000 cells in 24-well plates. Next day, parental:T1-160 mixtures were treated with 160 nM while parental:T2-450 mixtures were treated with 450 nM Taxol. Phase contrast, GFP and Texas Red images were taken as 2x2 with 10x objective using Cytation5 (BioTek, USA) in every 15 minutes for 72 hours. Number of mCherry+ and eGFP+ cells were counted from images using Gen5 software (BioTek, USA).

**Copy Number Variation Analysis.** Genomic DNAs of parental and resistant cell lines were isolated by using MN Nucleospin Tissue kit according to manufacturer's recommendation. 10 ng of genomic DNA was used as template for qPCR using SYBR Green, as per standard procedures. Primers used in copy number variation analysis are listed in **Supplementary Table 6.**

**Calcein Assay.** Calcein-AM is a membrane permeable dye that is converted to a fluorescent calcein by intracellular esterases. It is also a substrate for P-Glycoprotein (Pgp, ABCB1). Increased Pgp expression enhances the efflux of Calcein-AM and decreases fluorescence signal. Cells were seeded as 65 000 cells/well in 24-well plate. Next day, cells were treated with 10 µM Verapamil for 30 minutes at 37°C. Calcein-AM (C1430, Invitrogen) was added in final concentration of 125 nM and cells were incubated for another 30 minutes at 37°C. Images were taken with phase contrast and GFP filters as 2x2 with 10x objective using Cytation5 (BioTek, USA). Quantification of green fluorescence was performed in Synergy H1 Plate Reader (BioTek, USA).

### SUPPLEMENTARY TABLES

**Supplementary Table 1:** Antibodies used in this study

| **Primary Antibody** | Brand, Cat no |
|---|---|
| Total PARP | Abcam, ab74290 |
| Cleaved PARP | Cell Signaling, 9541 |
| Cleaved Caspase 3 | Cell Signaling, 9664 |
| GAPDH | Abcam, ab9485 |
| ABCB1 | Cell Signaling, 13978 |
| Tubulin | Abcam, ab15246 |
| BRPF1 | Novus, NBP2-15620 |
| H3 | Abcam, ab9049 |
| H3K27Ac | Cell Signaling, 4353 |
| H3K23Ac | Millipore, 07-355 |
| H3K14Ac | Cell Signaling, 7627 |
| H3K9Ac | Cell Signaling, 9649 |
| Puromycin | Merck Millipore, MABE342 |
| **Secondary Antibody** | **Brand, Cat no** |
| Goat anti-rabbit | Abcam, ab97051 |

**Supplementary Table 2:** qPCR primers used in this study

| Gene | Forward Primer (5' to 3') | Reverse Primer (5' to 3') |
|---|---|---|
| GAPDH | AGCCACATCGCTCAGACAC | GCCCAATACGACCAAATCC |
| BRPF1 | CCACACTGAAGATGCAGCCGA | CCGCAGTGCCGTCATCTCTT |
| BRPF2 | CAGCGGAAGAAGCAGTTTGTG | TCTTTGGCAGCCTTCATCTCC |
| BRPF3 | AGAACATCGGCTATGACCCC | CACCTCTGGGGACAAATGGG |
| RUNX1 | GGTTTCGCAGCGTGGTAAAA | GCACTGTGGGTACGAAGGAA |
| RUNX2 | CCGAGACCAACAGAGTCATTTA | ACATGGTGTCACTGTGCTGAA |
| JADE3 | AGACACCGTTCCACAGCCTTCT | ATGTAGCCAGGCTCTGTGGTGT |
| MEAF6 | GCGGCTCTTCAGTAAATCCTCG | GGAGAAGTGTCACTTTCCGTCC |
| ING5 | CAGAACGCCTACAGCAAGTGCA | CAGGTCTGCATCAAGCCTTCGA |
| KAT6A | CTTCAGTGAGAGCAGCGAGGAG | GTGGTGTTTGCGCTTTCGGACT |
| KAT6B | GGCAAGGATTTGGACGGTTTCTC | CGCTCTTCCAATATGCCAGGTAG |
| KAT7 | TCCATCTCAGGATGCCCACTGT | GTCATCTTGCCTGTGAGACAGC |
| ABCB1 | ACAGAGGGGATGGTCAGTGT | TCACGGCCATAGCGAATGTT |
| ABCC1 | CCGCTCTGGGACTGGAATG | ATGTAGCCTCGGTCATGTCG |
| ABCC3 | TATGCCCCCGATGAGGACCA | GACAGGGCACTCAGCTGTCTCA |

**Supplementary Table 3:** Composition of Chemical Probe Library and working concentrations of drugs in screens

| No | Source Name | Class/Target | Working Concentration (µM) |
|---|---|---|---|
| 1 | AMI-1 | Arginine methyltransferase - PRMT | 50 |
| 2 | SGC707 | Arginine methyltransferase - PRMT3 | 1 |
| 3 | TP-064 | Arginine methyltransferase - PRMT4 | 1 |
| 4 | TP-064N | Arginine methyltransferase - PRMT4 | 1 |
| 5 | MS049 | Arginine methyltransferase - PRMT4, PRMT6 | 1 |
| 6 | MS409N | Arginine methyltransferase - PRMT4, PRMT6 | 1 |
| 7 | GSK591 | Arginine methyltransferase - PRMT5 | 1 |
| 8 | LLY-283 | Arginine methyltransferase - PRMT5 | 1 |
| 9 | MS023 | Arginine methyltransferase - Type I PRMTs | 1 |
| 10 | GSK8814 | Bromodomains - ATAD2 | 10 |
| 11 | GSK8815 | Bromodomains - ATAD2 | 10 |
| 12 | GSK2801 | Bromodomains - BAZ2A, BAZ2B | 1 |
| 13 | BAZ2-ICR | Bromodomains - BAZ2A, BAZ2B | 1 |
| 14 | BAY-299 | Bromodomains - BRD1, TAF1 | 1 |
| 15 | RVX-208 | Bromodomains - BRD2, BRD3, BRD4, BRDT (BET, BD2) | 5 |
| 16 | (+)-JQ1 | Bromodomains - BRD2, BRD3, BRD4, BRDT (BET) | 1 |
| 17 | PFI-1 | Bromodomains - BRD2, BRD3, BRD4, BRDT (BET) | 5 |
| 18 | I-BET | Bromodomains - BRD2/3/4 | 1 |
| 19 | I-BRD9 | Bromodomains - BRD9 | 10 |
| 20 | TP-472 | Bromodomains - BRD9 | 1 |
| 21 | TP-472N | Bromodomains - BRD9 | 1 |
| 22 | LP99 | Bromodomains - BRD9, BRD7 | 1 |
| 23 | BI-9564 | Bromodomains - BRD9, BRD7 | 1 |
| 24 | GSK6853 | Bromodomains - BRPF1/2/3 | 1 |
| 25 | GSK9311 | Bromodomains - BRPF1/2/3 | 1 |
| 26 | PFI-4 | Bromodomains - BRPF1B | 1 |
| 27 | CBP/BRD4 (0383) | Bromodomains - CBP, BRD4(1) | 5 |
| 28 | NVS-CECR2-1 | Bromodomains - CECR2 | 1 |
| 29 | NVS-CECR2-C | Bromodomains - CECR2 | 1 |
| 30 | I-CBP112 | Bromodomains - CREBBP, EP300 | 1 |
| 31 | SGC-CBP30 | Bromodomains - CREBBP, EP300 | 1 |
| 32 | (-)-JQ1 (inactive) | Bromodomains - Negative control | 1 |
| 33 | Bromosporine | Bromodomains - pan-Bromodomain | 1 |
| 34 | OF-1 | Bromodomains - pan-BRPF | 5 |
| 35 | NI-57 | Bromodomains - pan-BRPF | 1 |
| 36 | GSK4027 | Bromodomains - PCAF, GCN5 | 1 |
| 37 | GSK4028 | Bromodomains - PCAF, GCN5 | 1 |
| 38 | L-Moses | Bromodomains - PCAF, GCN5 | 1 |
| 39 | D-Moses | Bromodomains - PCAF, GCN5 | 1 |
| 40 | SMARCA | Bromodomains - SMARCA, PB1 | 2.5 |
| 41 | PB1/SMARCA | Bromodomains - SMARCA, PB1 | 1 |
| 42 | PFI-3 | Bromodomains - SMARCA2/4, PB1(5) | 1 |

| | | | |
|---|---|---|---|
| 43 | TRIM24/BRPF | Bromodomains - TRIM24/BRPF | 10 |
| 44 | GSK864 | Dehydrogenase | 5 |
| 45 | 5-Azacitidine | DNA methyltransferase (DNMT) | 10 |
| 46 | 5-Azadeoxycitidine | DNA methyltransferase (DNMT) - DNMT1/3 | 5 |
| 47 | CXD101 | HDAC | 1 |
| 48 | PCI-24781 | HDAC | 10 |
| 49 | Romidepsin | HDAC | 1 |
| 50 | Mocetinostat | HDAC | 10 |
| 51 | CI-994 | HDAC - 1,2,3,(8) | 1 |
| 52 | Valproic acid | HDAC - aliphatic acid compounds | 1000 |
| 53 | RGFP966 | HDAC - HDAC3 | 10 |
| 54 | Rocilinostat | HDAC - HDAC6 | 10 |
| 55 | Tubastatin A HCl | HDAC - HDAC6 | 10 |
| 56 | PCI-34051 | HDAC - HDAC8 | 5 |
| 57 | Belinostat | HDAC - hydroxamic acids | 5 |
| 58 | SAHA | HDAC - hydroxamic acids | 2.5 |
| 59 | Trichostatin A | HDAC - hydroxamic acids - Class I & II | 0.5 |
| 60 | Entinostat | HDAC - ortho-amino anilides | 0.5 |
| 61 | EX 527 | HDAC - SIRT1 | 1 |
| 62 | SRT1720 | HDAC - SIRT1 (indirect) activator | 1 |
| 63 | AGK2 | HDAC - SIRT2 | 10 |
| 64 | TMP269 | HDAC -4, 5, 7 &9 | 10 |
| 65 | TMP195 | HDAC -4,5,7,9 | 1 |
| 66 | Santacruzamate | HDAC 2 | 50 |
| 67 | C646 | Histone acetyltransferase (HAT) p300/CBP | 1 |
| 68 | A-485 | Histone acetyltransferase (HAT) p300/CBP | 1 |
| 69 | A-486 | Histone acetyltransferase (HAT) p300/CBP | 1 |
| 70 | Methylstat (Ester) | Histone demethylase | 2.5 |
| 71 | KDOBA67 | Histone demethylase | 10 |
| 72 | KDM5-C70 | Histone demethylase - JARID1 | 10 |
| 73 | ML324 | Histone demethylase - JMJD2E | 5 |
| 74 | (E)-JIB-04 | Histone demethylase - Pan JmjC | 0.05 |
| 75 | SGC0946 | Histone methyltransferase - DOT1L | 7.5 |
| 76 | GSK343 | Histone methyltransferase - EZH2 | 3 |
| 77 | UNC1999 | Histone methyltransferase - EZH2 | 1 |
| 78 | UNC2400 | Histone methyltransferase - EZH2 | 1 |
| 79 | CPI-360 | Histone methyltransferase - EZH2 and EZH1 | 10 |
| 80 | CPI-169 | Histone methyltransferase - EZH2, EZH1 | 10 |
| 81 | UNC0642 | Histone methyltransferase - G9a, GLP | 1 |
| 82 | UNC0638 | Histone methyltransferase - G9a, GLP | 1 |
| 83 | A-366 | Histone methyltransferase - G9a, GLP | 2 |
| 84 | PFI-2 | Histone methyltransferase - SETD7 | 2 |
| 85 | LLY-507 | Histone methyltransferase - SMYD2 | 1 |
| 86 | BAY-598 | Histone methyltransferase - SMYD2 | 1 |
| 87 | PFI-5 | Histone methyltransferase - SMYD2 | 1 |
| 88 | Chaetocin | Histone methyltransferase - SUV39H1 | 0.05 |
| 89 | A-196 | Histone methyltransferase - SUV420H1/H2 | 1 |

| | | | |
|---|---|---|---|
| 90 | K00135 | Kinase inhibitor - ATP competitive - PIM | 1 |
| 91 | 5-Iodotubercidin | Kinase inhibitor - ATP mimetic - Haspin | 1 |
| 92 | SGI-1776 | Kinase inhibitor - Haspin | 10 |
| 93 | CHR-6494 | Kinase inhibitor - Haspin | 1 |
| 94 | KDOAM-25a | Lysine demethylases - JARID | 1 |
| 95 | KDOAM32 | Lysine demethylases - JARID | 1 |
| 96 | GSK J4 | Lysine demethylases - JMJD3, UTX, JARID1B | 10 |
| 97 | KDOPZ-32a | Lysine demethylases - KDM5 | 1 |
| 98 | Tranylcypromine | Lysine demethylases - LSD1 | 20 |
| 99 | GSK-LSD1 | Lysine demethylases - LSD1 - irreversible | 0.5 |
| 100 | GSK2879552 | Lysine demethylases - LSD1 | 10 |
| 101 | GSK J5 (inactive) | Lysine demethylases - Negative control | 10 |
| 102 | IOX1 | Lysine demethylases - pan-2-OG - (5-carboxy-8HQ) | 40 |
| 103 | KDOOA012000 | Lysine demethylases KDM2 | 1 |
| 104 | A-395 | Methyl Lysine Binder - EED | 1 |
| 105 | A-395N | Methyl Lysine Binder - EED | 1 |
| 106 | UNC1215 | Methyl Lysine Binder - L3MBTL3 | 5 |
| 107 | OICR-9429 | Methyl Lysine Binder - WDR5 | 1 |
| 108 | TDO20824a | Methyl Lysine Binder/tudor domain - Spin1 | 1 |
| 109 | TDO20826a | Methyl Lysine Binder/tudor domain - Spin1 | 1 |
| 110 | GSK484 | Peptidyl arginine deiminase (PAD4) | 1 |
| 111 | GSK106 | Peptidyl arginine deiminase (PAD4) | 1 |
| 112 | Olaparib | Poly ADP ribose polymerase (PARP) | 1 |
| 113 | Rucaparib | Poly ADP ribose polymerase (PARP) | 10 |
| 114 | IOX2 | Prolyl-Hydroxylases - PHD2 (EGLN1) | 10 |
| 115 | MAZ1805 | | 1 |
| 116 | MAZ1392 | | 1 |
| 117 | Bortezomib | | 0.1 |
| 118 | Carfilzomib | | 0.1 |

**Supplementary Table 4:** Sequences of sgRNAs used in this study

| Gene | Forward Sequence (5' to 3') | Reverse Sequence (5' to 3') |
|---|---|---|
| Non-targeting (NT1) | GACGGAGGCTAAGCGTCGCAA | TTGCGACGCTTAGCCTCCGTC |
| ABCB1_g1 | TTGGACTGTCAGCTGCTGTC | GACAGCAGCTGACAGTCCAA |
| BRPF1_g1 | AGGGTGACTGCAGGCAACGG | CCGTTGCCTGCAGTCACCCT |
| BRPF1_g2 | CCAACCGCCTGACCATCCAA | TTGGATGGTCAGGCGGTTGG |
| BRPF1_g3 | TGAGTACCTAATGGACCGAC | GTCGGTCCATTAGGTACTCA |
| BRPF2_g1 | CGACTCACCGGCTGCGATCC | GGATCGCAGCCGGTGAGTCG |
| BRPF2_g2 | GCAGCAGTCTCTGATCGACG | CGTCGATCAGAGACTGCTGC |
| BRPF3_g1 | AGAACCAGTCAACTTGAGTG | CACTCAAGTTGACTGGTTCT |
| BRPF3_g2 | GAGCGCCATGCGGTCCAGTG | CACTGGACCGCATGGCGCTC |
| KAT6A_g1 | TTCACTCGAACCGTTAGTTC | GAACTAACGGTTCGAGTGAA |
| KAT6B_g1 | TGAAAGACGGACCGCAGTAC | GTACTGCGGTCCGTCTTTCA |
| MEAF6_g1 | ATGTATGGCAATATTATTCG | CGAATAATATTGCCATACAT |
| ING5_g1 | CTTCCAGCTGATGCGAGAGC | GCTCTCGCATCAGCTGGAAG |
| JADE3_g1 | TCAGCATTGCTTGTCCTGAG | CTCAGGACAAGCAATGCTGA |
| KAT7_g1 | TCTCATCGTGAGATACATTG | CAATGTATCTCACGATGAGA |
| ARID1A_g1 | AATACTCACAGGCAAGCTGG | CCAGCTTGCCTGTGAGTATT |
| ARID1A_g2 | ATGGTCATCGGGTACCGCTG | CAGCGGTACCCGATGACCAT |
| GATAD1_g1 | AGAGTAAGCAGGAAATTCAC | GTGAATTTCCTGCTTACTCT |
| GATAD1_g2 | CGTGACTTGAAATACTCAGA | TCTGAGTATTTCAAGTCACG |
| SMARCE1_g1 | ACCAACAGCCGGGTCACGGT | ACCGTGACCCGGCTGTTGGT |
| SMARCE1_g2 | TATGTAAGCAAGGTACGCGG | CCGCGTACCTTGCTTACATA |
| KMT2A_g1 | AGAAAGGACGTCGATCGAGG | CCTCGATCGACGTCCTTTCT |
| KMT2A_g2 | TCAGAGTGCGAAGTCCCACA | TGTGGGACTTCGCACTCTGA |
| MEN1_g1 | CATGCGCTGTGACCGCAAGA | TCTTGCGGTCACAGCGCATG |
| MEN1_g2 | CCAGGCATGATCCTCAGACA | TGTCTGAGGATCATGCCTGG |
| ARNTL_g1 | CTGGACATTGCGTTGCATGT | ACATGCAACGCAATGTCCAG |
| ARNTL_g2 | TTAGAATATACAGAACACCA | TGGTGTTCTGTATATTCTAA |
| BRD8_g1 | AGGAGGTGATTATCCACTTG | CAAGTGGATAATCACCTCCT |
| BRD8_g2 | ATAAGTACCTATATCTCTCC | GGAGAGATATAGGTACTTAT |
| MLLT6_g1 | AATCTCAGGAGCGAGCAGCC | GGCTGCTCGCTCCTGAGATT |
| MLLT6_g2 | AGCTTGCTATGGCATCGTTC | GAACGATGCCATAGCAAGCT |
| BRD3_g1 | CGACGTGACGTTTGCAGTGA | TCACTGCAAACGTCACGTCG |
| BRD3_g2 | GAGGAGAGCTCTTCGGACTC | GAGTCCGAAGAGCTCTCCTC |
| PPP2CA_g1 | AATAAAAGTCATACCTCATG | CATGAGGTATGACTTTTATT |
| PPP2CA_g2 | GGTATATCTCCTCGAGGAGC | GCTCCTCGAGGAGATATACC |
| CHD8_g1 | CTGTCTTCTACACTACCGTG | CACGGTAGTGTAGAAGACAG |
| CHD8_g2 | CTTTAATCCAGACTACGTAG | CTACGTAGTCTGGATTAAAG |

**Supplementary Table 5:** ChIP-QPCR primers used in this study

| Gene | Forward Primer (5' to 3') | Reverse Primer (5' to 3') |
|---|---|---|
| ABCB1 promoter (-658 bp) | TTCTCTCTGTGACAGCTCAGT | AGCACAAATTGAAGGAAGGAGT |
| RUNX2 Acetylation | GGAAACGGAGGGGTCTGAAC | TAAGAGCTGGTTTCGCCGTC |
| Chr 12 Gene Desert (negative cnt) | TGTACAGGGCCTGGTTACCCACA | AGGAGGGCCTAGCTGGTGTCAT |

**Supplementary Table 6:** q-RT-PCR from genomic DNA primers used in this study

| Gene | Forward Primer (5' to 3') | Reverse Primer (5' to 3') |
|---|---|---|
| GAPDH | CGGCTACTAGCGGTTTTACG | AAGAAGATGCGGCTGACTGT |
| ABCB1 | AGATCTACCAGGACGAGTGAGAAAA | AACAGTCAGTTCCTATATCCTGTGTCT |

### SUPPLEMENTARY FIGURES

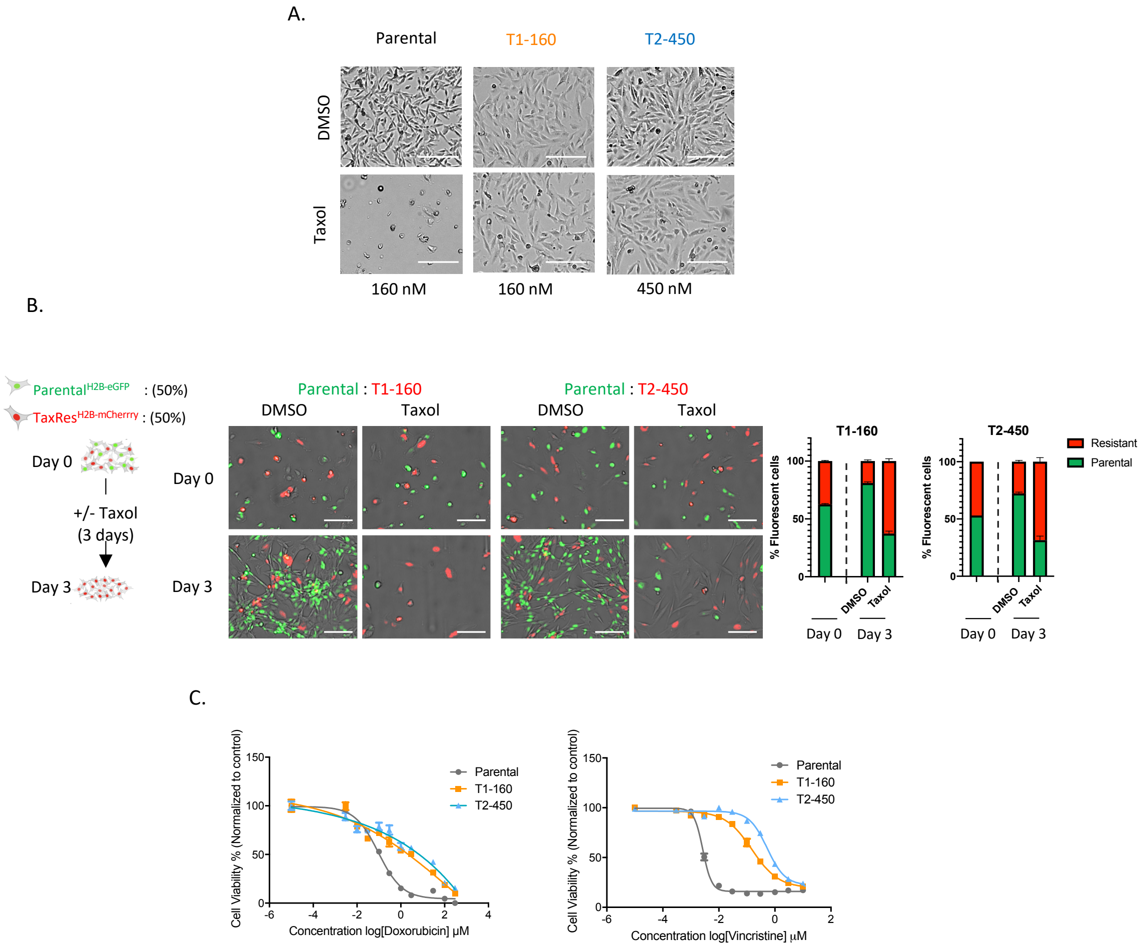


**Supplementary Figure 1. Taxol-resistant phenotype. A.** Representative images from live cell imaging. Cells were seeded to 12-well plates in the presence or absence of Taxol (Parental: 160 nM, T1-160: 160 nM, T2-450: 450 nM) and images were taken at every 15 minutes for 3 days after drug treatment. Scale bar: 200 µm**. B.** Competition assay in the presence of Taxol. Parental cells were labeled with PGK-H2B-eGFP virus, resistant cells were labeled with pGK-H2B-mCherry and mixed in 1:1 ratio. After +/- Taxol treatment, images were taken at every 20 minutes for 3 days after drug treatment. Scale bar: 100 µm. **C.** Cross-resistance phenotype of T1-160 and T2-450 cells upon Doxorubicin and Vincristine treatment.

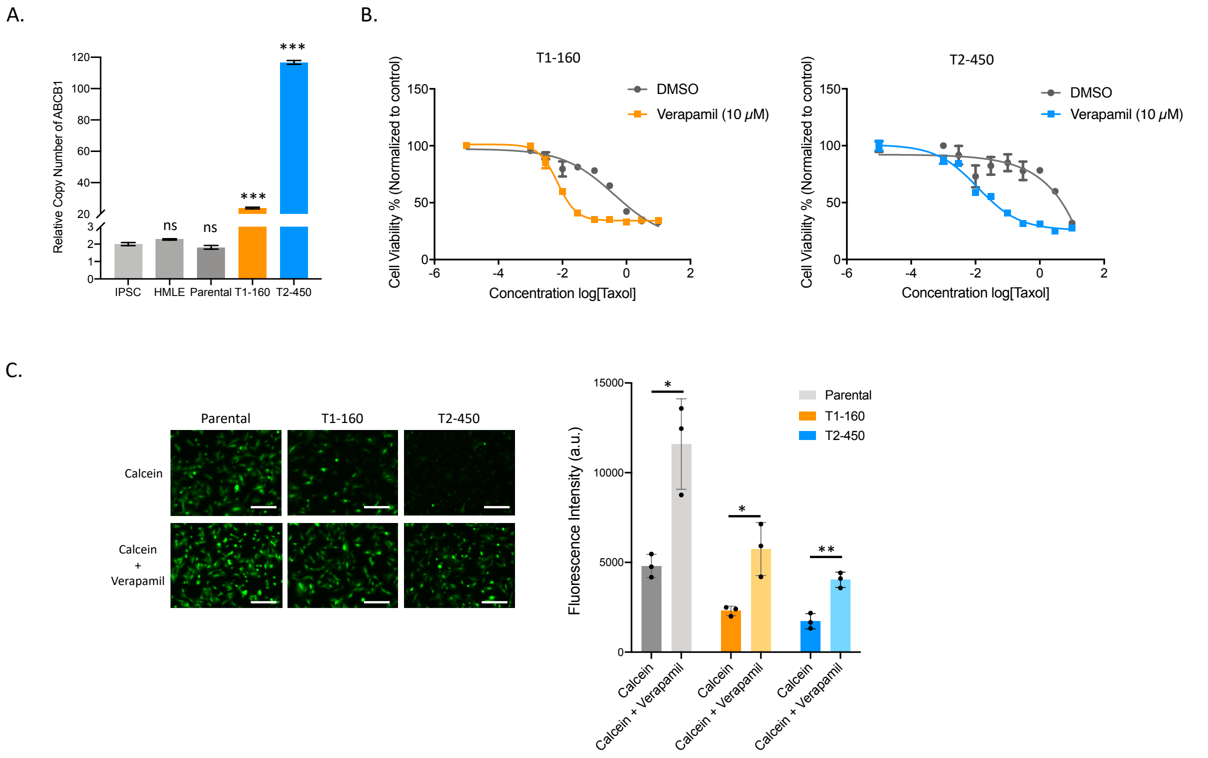


**Supplementary Figure 2. ABCB1 expression and function in Taxol-resistant cells. A.** Copy number variation (CNV) analysis of ABCB1 in Taxol-resistant cells when compared to control cell lines (HMLE and IPSC) with normal karyotypes. **B.** Effect of verapamil on cell viability of Taxol-resistant T1-160 (left) and T2-450 cells (right). **C.** Representative images of Calcein-AM (125 nM) assay in the presence of verapamil (10 μM) on parental and resistant cells. Scale bar: 100 μm. Quantification of fluorescent signal was performed using microplate reader.

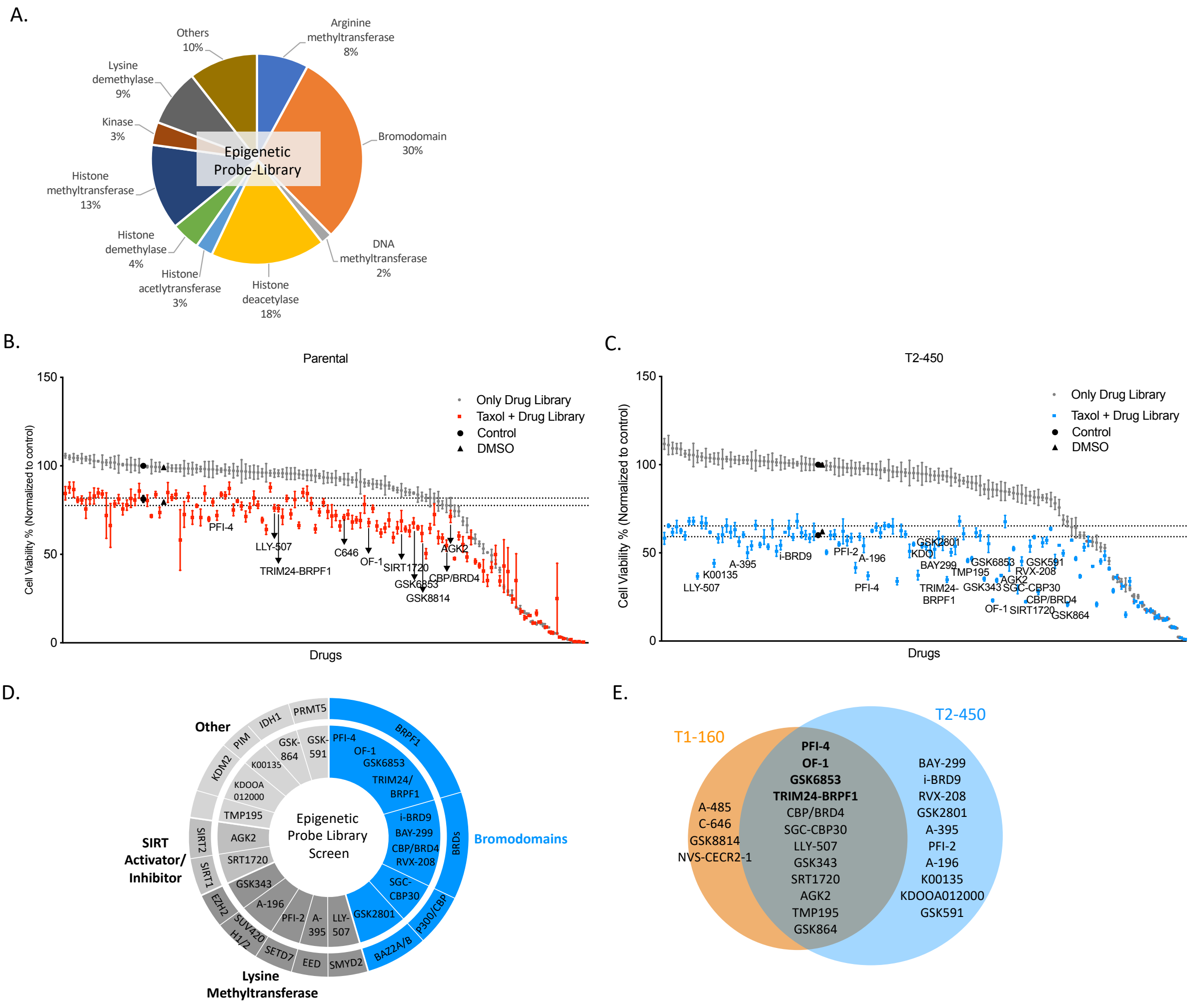


**Supplementary Figure 3. Epigenetic probe library screen on parental and T2-450 cells. A.** Pie chart shows the epigenetic modifier classes targeted by the epigenetic chemical probe library. **B.** Results of epigenetic probe library screen performed on SUM159PT parental cells. **C.** Results of epigenetic probe library screen performed on T2-450 cells. Both in (B.) and (C.) grey dots represent the effect of epigenetic probes alone while the colored dots show changes in cell viability when Taxol was combined with a specific epigenetic probe. Epigenetic probes that significantly reversed drug resistance are labeled on the T2-450 cell graph. If the negative control of a given probe also showed a significant depletion in cell viability, neither of the probes was labeled on the graph. In Parental cell screen results, the epigenetic probes identified as sensitizers in Taxol-resistant cells were labeled **D.** Classification of hits identified in epigenetic probe library screen of T2-450 cells. Target gene of the chemical probes were indicated in the outer circle. **E.** Venn diagram illustrates the common probes showing significant cell viability decrease in T1-160 and T2-450 epigenetic probe library screens.

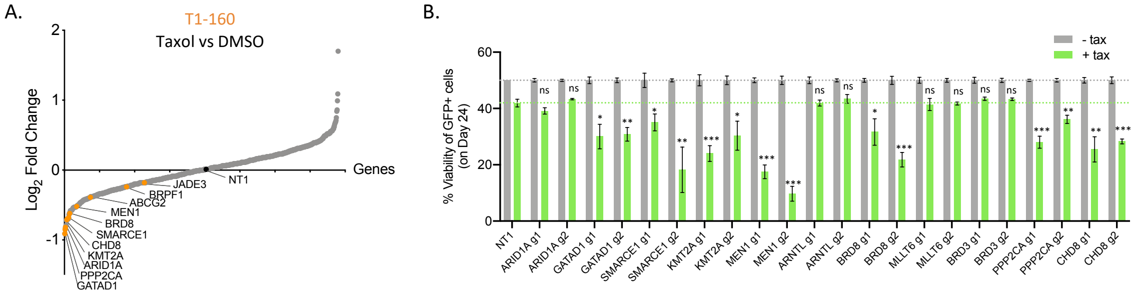


**Supplementary Figure 4. Competition assay results of EPIKOL screen on T1-160 cells A.** Waterfall plot of EPIKOL screen results showing $Log_2$FoldChanges of genes used in dual-color competition assay **B.** Dual-color competition assay results of candidate genes from T1-160 EPIKOL screen. Gray bars represent the eGFP+ cells in eGFP:mCherry mixtures in the absence of Taxol. Green bars indicate the percentage of eGFP+ cells carrying sgRNA of interest in the population in the presence of Taxol. Untreated sgRNA mixtures were normalized to 50%. P values determined by two-tailed Student's t-test in comparison to control group; ns >0.05, *p < 0.05.

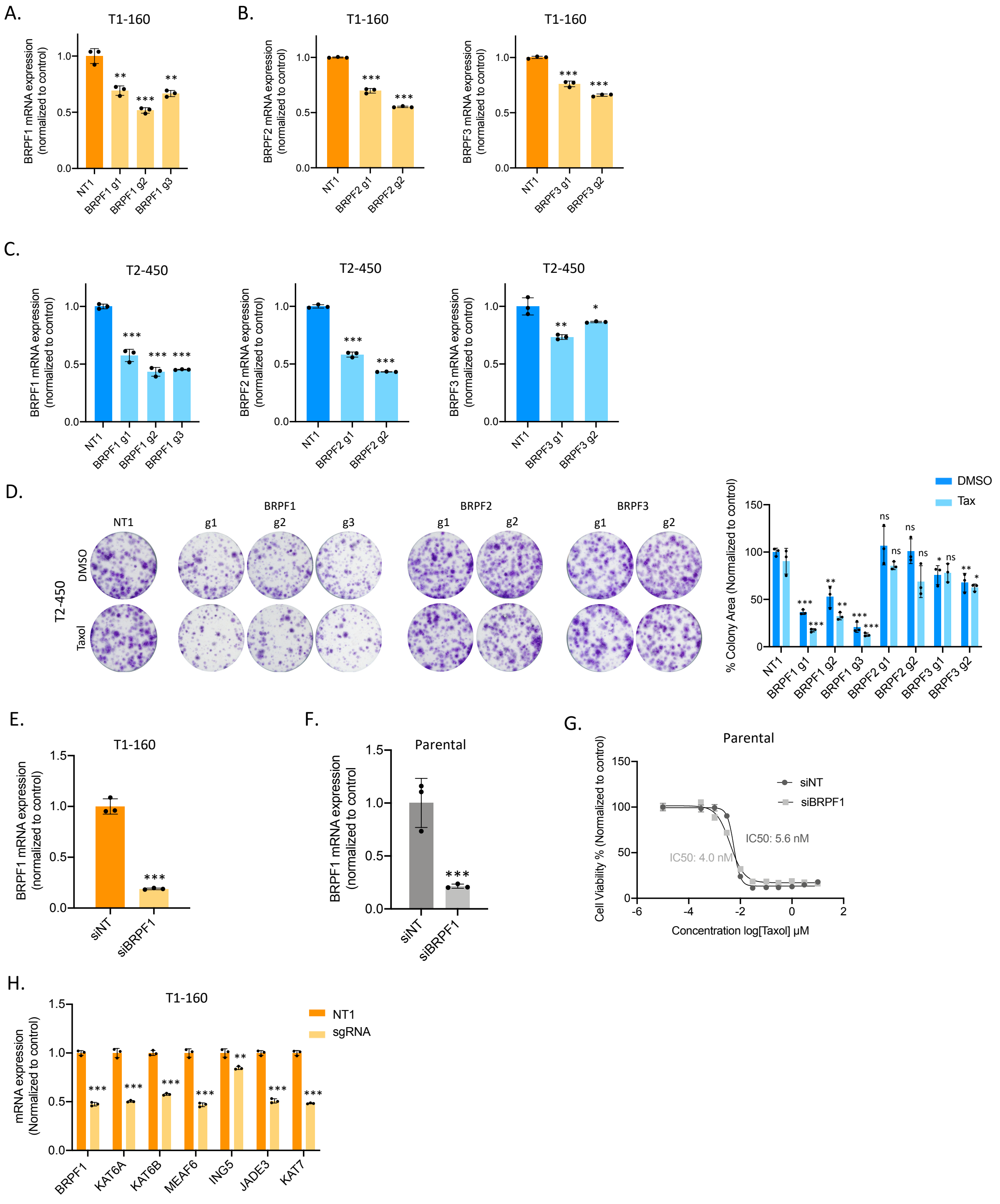


**Supplementary Figure 5. Effects of BRPF1, BRPF2, BRPF3 knockout on Parental and Taxol-resistant cells. A.** BRPF1 expression in mRNA level upon BRPF1 knockout in T1-160 cells. **B.** BRPF2 and BRPF3 mRNA expression levels upon their knockouts in T1-160 cells. **C.** BRPF1, BRPF2 and BRPF3 mRNA expression levels upon their knockouts on T2-450 cells **D.** Clonogenic assay showing the effect of BRPF1, BRPF2, and BRPF3 knockouts in the presence of Taxol and quantification of the colonies. **E.** BRPF1 mRNA levels upon BRPF1 siRNA transfection in T1-160 cells. **F.** BRPF1 mRNA levels upon BRPF1 siRNA transfection in Parental cells. **G.** Cell viability measurement performed with siNT and siBRPF1 samples in the presence of Taxol on Parental cells. **H.** mRNA expression levels of MOZ/MORF and HBO1 complex members upon knockout in T1-160 cells. P values determined by two-tailed Student's t-test in comparison to control group; *p < 0.05, **p < 0.01, ***p < 0.001.

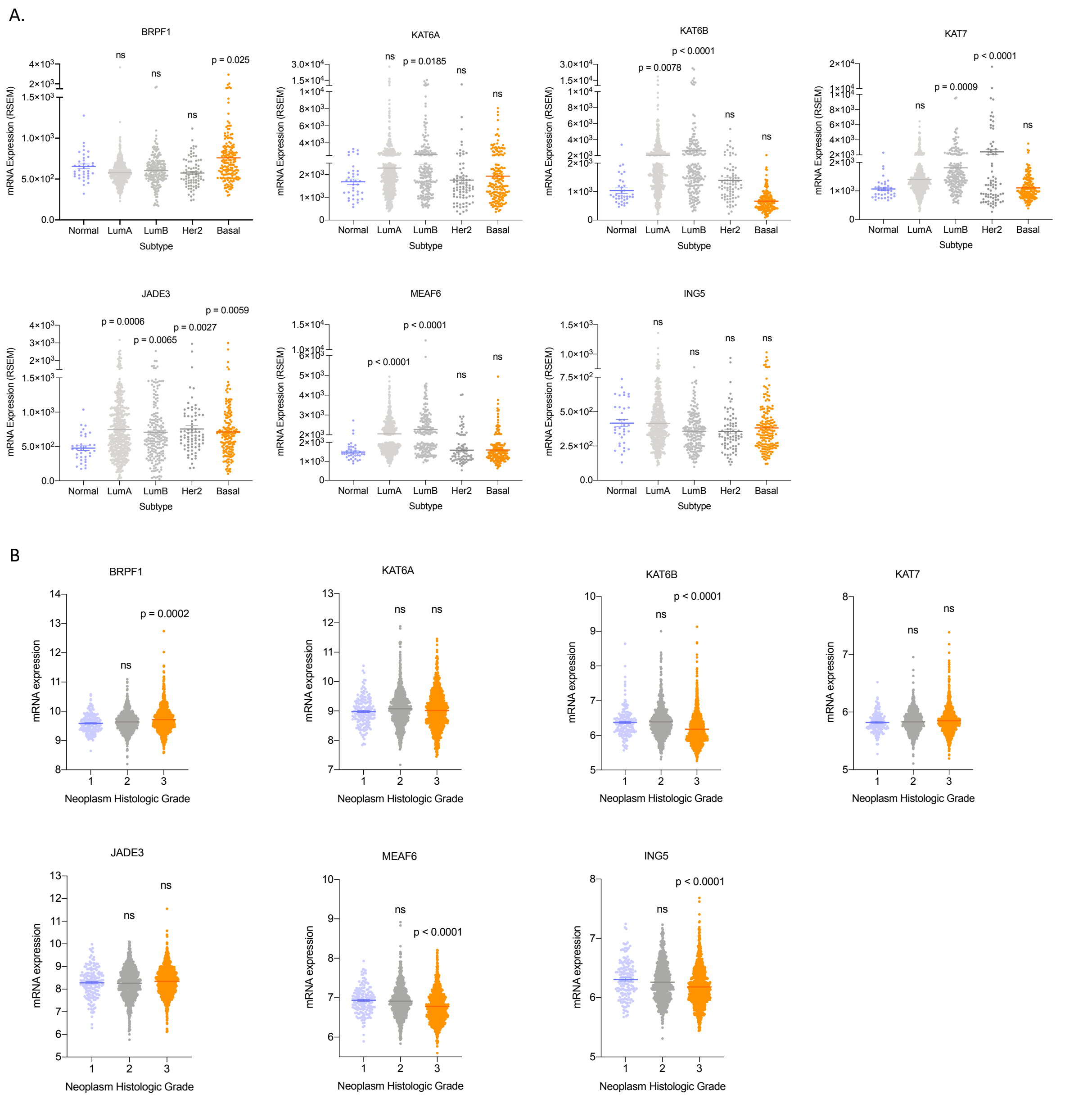


**Supplementary Figure 6. MOZ/MORF and HBO1 complex members' expression analysis in breast cancer subtypes and histological grades.** mRNA levels of MOZ/MORF and HBO1 complex members in clinical samples were accessed through cBioPortal. **A.** Relationship between the expression of the gene and subtype was analyzed using TCGA - Breast Invasive Carcinoma - PanCancer Atlas. Ordinary one-way Anova was performed by comparing the groups to Normal subtype and p values were written above the group if it is $p < 0.05$ **B.** Histological grade analysis was performed using METABRIC database of breast cancer. Ordinary one-way Anova was performed by comparing the higher histological grades to grade 1 and p values were written above the group if it is $p < 0.05$.

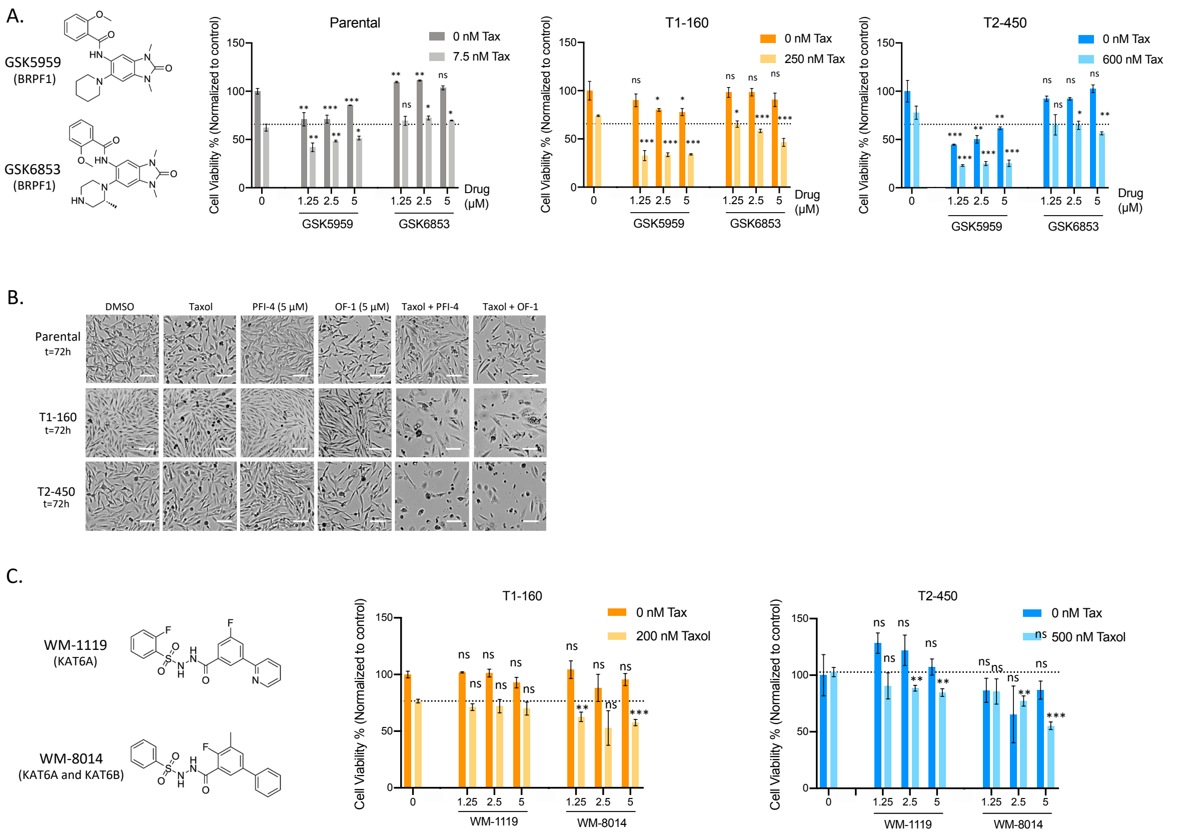

**Supplementary Figure 7. BRPF1 and MOZ/MORF inhibitor effects on parental and Taxol-resistant cells. A.** (Left) Chemical structures of BRPF1 inhibitors GSK5959 and GSK6853. GSK5959 specifically targets BRPF-1 but it was not present in epigenetic probe library. GSK6853 is another BRPF1 inhibitor identified during epigenetic probe library screens in both Taxol resistant cells. (Right) Cell viability results of combination of GSK5959 and GSK6853 with Taxol on Parental and Taxol-resistant cells **B.** Representative images from live-cell imaging in the presence of PFI-4, OF-1 and Taxol on Parental and Taxol-resistant cells. Scale bar: 100 µm **C.** (Left) Chemical structures of KAT6A/B inhibitors and (Right) cell viability results of their combination with Taxol in T1-160 and T2-450 cells. P values determined by two-tailed Student's t-test in comparison to control group; *p < 0.05, **p < 0.01, ***p < 0.001.

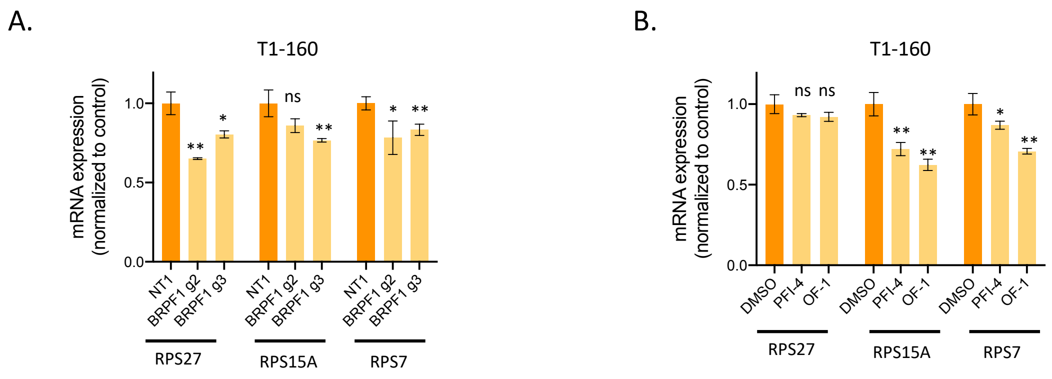


**Supplementary Figure 8. mRNA expression of selected ribosome-related genes upon BRPF1 loss or inhibition. A.** Ribosome-related gene mRNA expressions in BRPF1 KO T1-160 cells. **B.** Ribosome-related gene mRNA expressions in BRPF1 inhibitor treated T1-160 cells. P values determined by two-tailed Student's t-test in comparison to control group; *p < 0.05, **p < 0.01, ***p < 0.001.

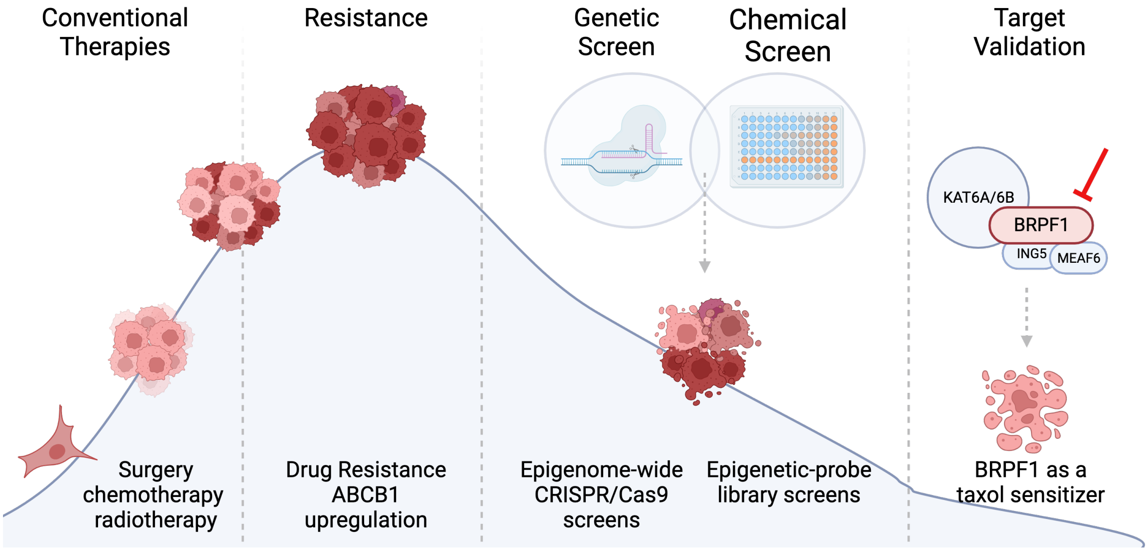


**Supplementary Figure 9. Graphical abstract.** Figure summarizes the approach of the study. Figure was created with BioRender.com.
